# SF3B3 / SF3B5 form a metazoan specific transcription module of the U2 snRNP that coordinates Pol II elongation in a splicing independent manner

**DOI:** 10.64898/2026.07.14.737342

**Authors:** Dane Vassiliadis, Jesse J. Balic, Olivia Braniff, Andrea Gillespie, William Rothnie, Kelsy Prest, Oliver Sinclair, Andrew Das, Ching-Seng Ang, Mark A. Dawson

**Author notes:** **Corresponding Authors:** Professor Mark Dawson, Peter MacCallum Cancer Centre, 305 Grattan Street, Melbourne VIC 3000, Australia, Doctor Dane Vassiliadis, Peter MacCallum Cancer Centre, 305 Grattan Street, Melbourne VIC 3000, Australia. These authors contributed equally to this work (Dane Vassiliadis, Jesse J. Balic).

## Abstract

Co-transcriptional splicing is a conserved feature of eukaryotic gene expression. However, establishing the functional nature of this process has been difficult. Here using high throughput CRISPR/Cas9 screens we surprisingly find that SF3B3, the third largest subunit of the U2 snRNP complex, is a major regulator of RNA Pol II pause release and processivity. Remarkably, the absence of SF3B3 dramatically perturbs transcription but U2 snRNP assembly and RNA splicing remains unaffected. Mechanistically, SF3B3 coordinates the chromatin occupancy of transcriptional kinases (CDK9/12/13) alongside the PAF1c and Integrator complexes to regulate Pol II. Structure / function analyses of SF3B3 revealed that a metazoan specific 18aa sequence within its disordered tail phenocopies its loss and mediates the physical association and stability of SF3B5. We show that loss of SF3B5 mirrors SF3B3 deficiency suggesting this submodule, although resident within the U2 snRNP complex, evolved to primarily coordinate RNA Pol II in a splicing-independent manner.

## Main

Transcription is a dynamic process whereby upstream signalling cascades act upon transcription factors (TFs), transcriptional co-activators and megadalton-scale protein complexes to coordinate the initiation, elongation and termination of RNA Polymerase II (Pol II). Co-activators such as the Mediator complex, the BET bromodomain family members BRD2/3/4 and the lysine acetyltransferases EP300/CREBBP (hereafter P300 and CBP) play important roles in gene regulation control^1–6^ and have emerged as key dependencies and therapeutic targets in cancer and other diseases^7–12^. Transcription is further nuanced by a physical and functional coupling to co-transcriptional processes including pre-mRNA splicing, RNA capping, 3’-end processing and polyadenylation^13–15^, which impart additional layers of regulation that refine gene expression through interactions between the RNA processing complexes and transcriptional machinery^16–18^.

It is well established that Pol II activity is enhanced by the presence of introns, a hallmark of eukaryotic protein-coding genes^19–22^. In mammals, introns make up ∼90% of the gene length and numerous studies have revealed that splicing is most efficient as a co-transcriptional process that occurs in the context of the chromatin template juxtaposed to Pol II and its associated elongation factors that regulate the rate of mRNA synthesis^23^. Central to the integration of the various inputs that facilitate co-transcriptional processes including splicing is the C-terminal domain (CTD) of Pol II, a tail-like structure comprised of 52 repeats of a hepta-peptide (YSPTSPS) sequence that is heavily modified by kinases and other enzymes^24–30^, and mediates interactions with a broad range of transcriptional complexes^25^. Truncation of the Pol II CTD markedly attenuates the efficiency of splicing^26,31^, whereas phosphorylation of the CTD potentiates splicing^32^. This reciprocal interplay is further supported by the fact that previous work has shown that splicing factors assemble onto the Pol II CTD region in a Serine 5 phosphorylation dependent manner proximal to nascent RNA^17,18,33,34^.

Whilst the physical interaction, including structural insights^35^, between the splicing and transcription machinery are well established and there is broad consensus that splicing is most efficient as a co-transcriptional process, mechanistic insights into how splicing factors influence the crosstalk between transcription and splicing remain unresolved. As with any coupled process, significant disruption to one component will negatively impact the other, making it challenging to separate correlation from causation. For instance, global inhibition of transcription with a CDK7 inhibitor causes a widespread disruption in splicing^36^. Similarly, inhibition of the U1 (by antisense morpholinos to U1 snRNA) or U2 (using Pladienolide B, an inhibitor of SF3B1) snRNP complexes markedly disrupt both splicing and transcription^37–39^. Therefore, it remains unclear if splicing factors have any direct influence on transcription or if the phenotypes observed by inhibiting these complexes are indirect and mediated by the uncoupling of a co-transcriptional activity. To gain further insights into the processes that govern the functional interaction between the core transcriptional apparatus, associated co-activators and RNA processing machinery we developed an approach that combines the real-time monitoring of endogenous gene expression with unbiased CRISPR screens and orthogonal degron based strategies. Together our findings uncover a critical splicing-independent role for the U2 spliceosome components SF3B3 and SF3B5 in modulating transcription.

### Endogenous reporter screens reveal that transcriptional co-activators have a subtle influence on gene expression

To develop a system that accurately monitors gene expression in live cells we used CRISPR/Cas9 to engineer an mNeonGreen (mNG, analogous to GFP) fluorescent reporter^40^ into the 3’ end of the endogenous *c-MYC* (hereafter *MYC*) locus in K562 cells (**Figure 1A**). We chose *MYC* as an exemplar gene to monitor endogenous transcriptional activity as it is a multi-intronic gene with a short mRNA and protein half-life and the network of transcriptional regulators controlling its expression have been studied in detail^41^. We isolated multiple single cell clones containing a homozygous integration of the mNG construct into the *MYC* locus (**Figure S1A**). These clones were positive for MYC-mNeonGreen (MYC-mNG) without any detrimental effects on cell proliferation (**Figures S1B-C**). Moreover, RNA-seq and cycloheximide chase assays confirmed that MYC-mNG cells were transcriptionally similar to parental K562 cells and had a comparable protein half-life to wild type MYC (**Figures S1D-E**).

**Figure 1.**
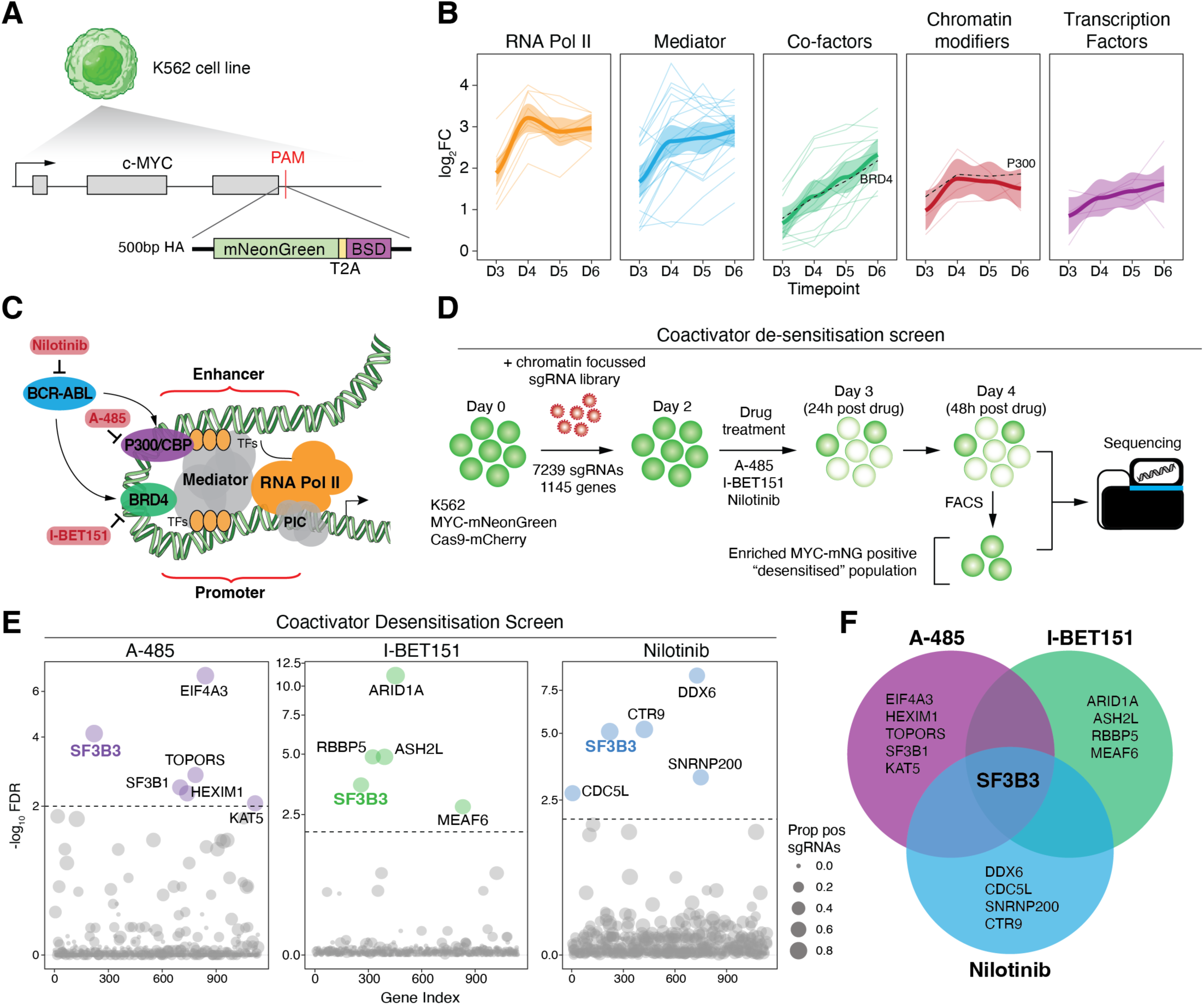
Loss of SF3B3 overcomes transcriptional co-activator inhibition. A) Schematic of mNeonGreen-T2A-Blasticidin cassette knock-in into the endogenous *c-MYC* locus in human K562 cells. B) Average enrichment profiles of core MYC regulators Pol II (orange) and Mediator (blue), or co-factors (green), chromatin modifiers (red) and transcription factors (purple) across days 3 to 6 in the temporal dependency screen. Mean log2 fold change (log_2_FC) versus library control (bold line) +/- standard deviation (confidence interval) are shown. Individual gene lineplots are shown in background. Results are the average of four independent biological replicates. C) Schematic of selected components of the basal transcriptional machinery, transcriptional co-activators and upstream signalling kinases that impinge on RNA Pol II during transcription. In red are small molecules that specifically inhibit the indicated factors. D) Schematic of the coactivator de-sensitisation CRISPR screen. E) Dotplots of results for A-485, I-BET151 and Nilotinib coactivator de-sensitisation screens. Significantly enriched hits are coloured. SF3B3 is shown in bold. Dashed lines indicate an adjusted p-value threshold of 0.01 (-log10 FDR = 2). F) Venn diagram showing overlap of enriched targets in all three coactivator de-sensitisation screens.

Having established an exemplar real-time readout for endogenous gene expression, we first studied the temporal nature by which gene expression is altered by performing a CRISPR/Cas9 “temporal dependency” screen targeting 1145 chromatin related factors (**Table S1, Figures S1F-G**). Our previous work studying transcriptional regulators (including common essential targets) with a CRISPR knockout strategy revealed that peak loss in gene expression is seen 4 days after guide infection^42,43^. This reflects the timeframe required for viral transgene integration into the host genome, guide RNA transcription, genome editing via Cas9, and finally, the turnover of pre-existing mRNA and protein. Therefore, to capture dynamic changes in reporter gene expression our screen captured four consecutive time points (Days 3, 4, 5 and 6) post-sgRNA transduction. This approach also caters for stochastic differences in sgRNA expression or editing efficiency^42^. The results clearly showed distinctive patterns of influence on reporter gene expression for different groups of chromatin factors in both kinetics and magnitude of effect (**Figure S1H**). As expected, knockout of components of the core transcriptional machinery including the RNA Polymerase II (Pol II) complex exhibited an early, profound and near uniform enrichment in the screen (**Figure 1B**). Similarly, components of the head, middle and tail modules of Mediator, which ensure the structural and functional integrity of the complex^44^, had a similar consequence to loss of the Pol II holoenzyme. Comparatively, knockout of transcription co-factors, including BRD4, displayed more gradual enrichment kinetics across the screen time course. Notably, most transcription factors and chromatin modifying enzymes, except for P300, showed weaker enrichment in the screen, likely reflecting both functional redundancy and/or an expansion of paralogs within mammals.

### The function of transcriptional co-activators is regulated by SF3B3

Targeting transcriptional coactivators has been clinically disappointing, due in part to the restoration of mRNA expression despite ongoing therapeutic pressure^45,46^. We therefore sought to discover previously unrecognized mediators that govern transcriptional repression following coactivator inhibition, as it may reveal novel insights into normal transcriptional regulation. To study this process, we targeted different factors that potentiate transcription (**Figure 1C**): (i) A-485, a catalytic inhibitor of CBP/P300^11^ (ii) I-BET151, a BET bromodomain inhibitor that targets BRD2, BRD3, BRD4 and BRDT^47^ and (iii) Nilotinib^48^, an ABL1 tyrosine kinase inhibitor that broadly inhibits the activity of several downstream transcription factors and associated co-factors required for gene expression in K562 cells. Each of these approaches resulted in a rapid decrease in our endogenous reporter of gene expression (**Figures S2A-B**) enabling a “coactivator de-sensitisation” CRISPR screen to identify common targets that overcome the inhibitory effects imposed on these various transcriptional regulators (**Figure 1D & Figure S2C**). Whilst several hits in our screen were unique dependencies to each of the cofactor inhibitors used, a notable and unexpected finding was that SF3B3, an evolutionarily conserved component of the U2 snRNP spliceosome complex, was a common dependency in all our screens (**Figures 1E-F**).

Using two independent sgRNAs targeting *SF3B3,* we validated the findings of our CRISPR screen (**Figure S2D**). To provide an orthogonal approach to the CRISPR knockout data, we next employed the dTAG inducible degron strategy to study the consequences of SF3B3 loss in the context of coactivator inhibition^49^. Here we endogenously tagged the C-terminus of *SF3B3* with the dTAG degron (FKBP12^F36V^), hereafter called SF3B3-dTAG (**Figure S2E**)^50,51^. We derived multiple clonal homozygous knock-in populations that showed near complete degradation of SF3B3 by 3 hours post dTAG^-V1^ treatment, whereas cells treated for up to 24 hours with dTAG^NEG^, an inactive dTAG^-V1^ enantiomer^52^, continue to express SF3B3 to wild type levels (**Figure S2F)**. We also confirmed by RNA-seq that these SF3B3-dTAG cells were transcriptionally equivalent to parental MYC-mNG cells at steady state (**Figure S2G**).

Consistent with the essential requirement for SF3B3 in eukaryotic cells, we observed a decrease in cell viability and reporter gene expression from 48-96 hours post dTAG^-V1^ treatment (**Figure S2H**). Consequently, we confined our degron-based assays to a maximum period of 24 hours of dTAG^-V1^ treatment, a timepoint at which we observed complete SF3B3 degradation (**Figure S2F**) with no discernible change in either viability or reporter expression via flow cytometry (**Figure S2H**). As CBP/P300 have long been considered the archetypal example of a transcriptional coactivator^53^ we used A-485 as an exemplar of coactivator inhibition. These data confirmed that cells with SF3B3 degraded phenocopy our results with genetic knockout of SF3B3 (**Figure 2A**). We also performed a complementary experiment whereby we first inhibited CBP/P300 and then degraded SF3B3 with dTAG^-V1^. Here, SF3B3 degradation also resulted in the recovery of reporter expression (**Figure S2I**). Next, we examined the kinetics of reporter gene expression following SF3B3 degradation. Interestingly, we found that expression was reduced to the same nadir and with the same kinetics in the presence or absence of SF3B3 suggesting that SF3B3 does not play a role in the active repression of transcription (**Figure S2J**). Notably, the repression following co-activator inhibition was not maintained in cells where SF3B3 is degraded. Therefore, in the absence of SF3B3, transcription can recommence despite the ongoing inhibition of critical co-activators such as the BET bromodomain proteins and CBP/P300 raising the possibility that SF3B3 provides key regulatory inputs that influence the activity of Pol II.

**Figure 2.**
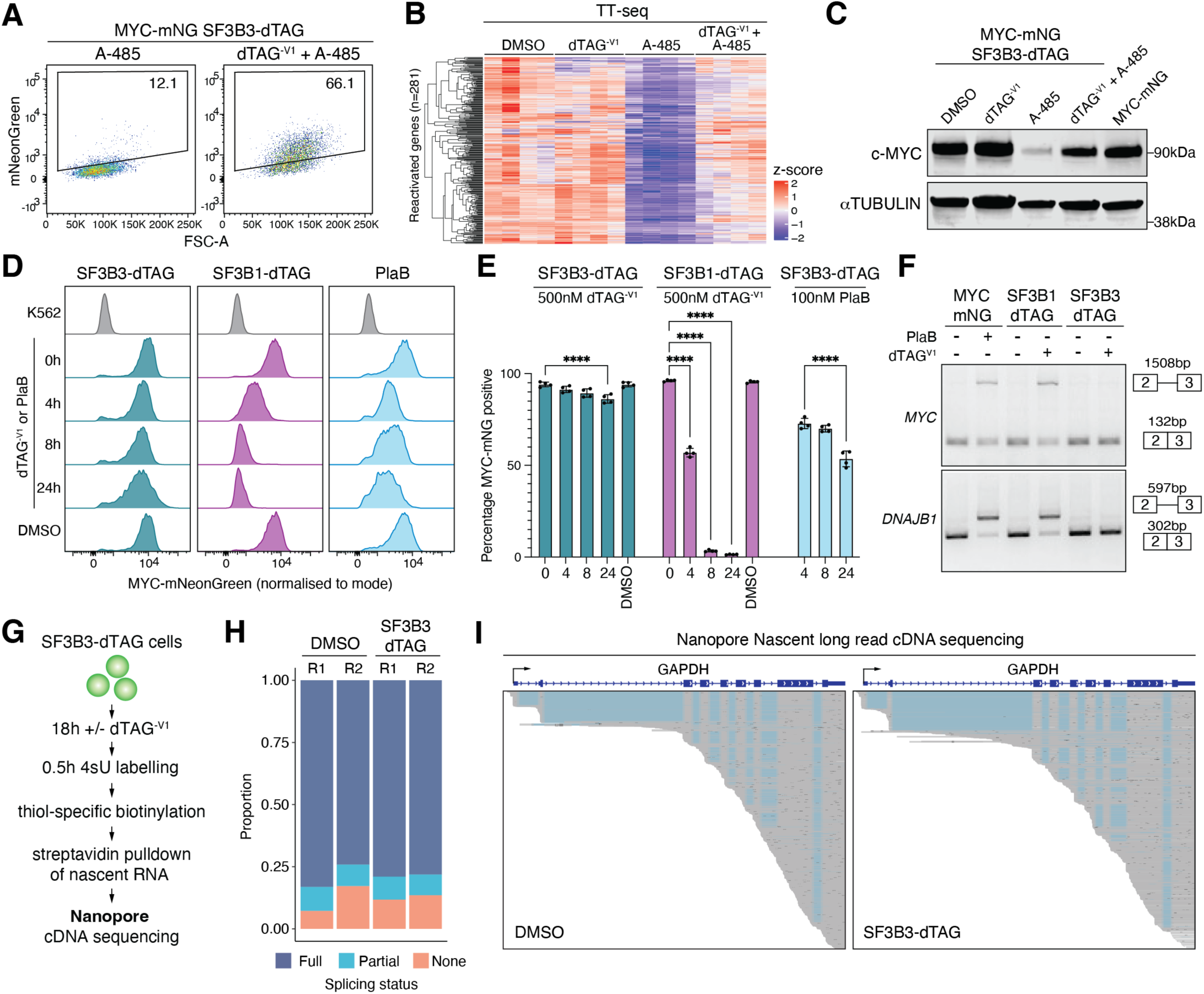
A splicing independent role for SF3B3 in transcriptional regulation. A) Flow cytometry dotplots of MYC-mNG expression in MYC-mNG SF3B3-dTAG cells pretreated for 3h with either 500nM dTAG^-V1^ or 500nM dTAG^-NEG^ followed by consecutive treatment with 2µM A-485 for a further 16h. Percentage of MYC-mNG positive cells are indicated. Representative of n = 3 independent experiments. B) Heatmap showing row-scaled TT-seq signal (n = 4 replicates) for reactivated genes (n = 281) in the SF3B3-dTAG + A-485 treatment condition. C) Western blot validation of MYC-mNG expression for the indicated treatments in MYC-mNG SF3B3-dTAG cells. Representative of three independent clones. MYC-mNG cells are included as a positive control. D) Flow cytometry histograms of MYC-mNG expression (normalised to mode) in SF3B3-dTAG (purple), SF3B1-dTAG (pink) or PlaB treated SF3B3-dTAG cells (blue). SF3B3-dTAG and SF3B1-dTAG cells were treated with 500nM dTAG^-V1^ for 0, 4, 8 and 24h. PlaB treated SF3B3-dTAG cells were treated with 100nM PlaB for 0, 4, 8 and 24h. DMSO indicates 24h treatment with 0.1% v/v DMSO. Representative of n = 4 independent experiments. E) Quantification of percentage MYC-mNG positive cells via flow cytometry data shown in (D). 0h and DMSO controls samples for SF3B3-dTAG cells treated with PlaB are analogous to the same samples in SF3B3-dTAG cells treated with dTAG^-V1^. Histograms show the mean and error bars show the standard deviation. n = 4 independent experiments. Two-sided T-test, **** = p < 0.001. F) Intron inclusion PCR assay across *c-MYC* exons 2-3 or *DNAJB1* exons 2-3 from MYC-mNG cells pretreated for 4 hours with 0.1% v/v DMSO or 100nM PlaB, or SF3B1-dTAG and SF3B3-dTAG cells pretreated for 4 hours with 0.1% v/v DMSO or 500nM dTAG^-V1^. Schematic indicates expected size of spliced and unspliced amplicons. G) Schematic of Oxford Nanopore experiment showing nascent RNA labelling, biotinylation and purification, reverse transcription and Nanopore cDNA sequencing. H) Classification of splicing status for Nanopore long read sequencing in DMSO or SF3B3-dTAG treatments. N = 2 independent replicates. Full = All introns covered by a read are spliced, Partial = some but not all introns covered by a read are spliced, None = No covered introns are spliced. I) Exemplar Nanopore long read alignment results at the human *GAPDH* locus for SF3B3-dTAG cells treated for 18h with 0.1 % v/v DMSO (DMSO) or 500nM dTAG^-V1^ (SF3B3-dTAG). Reads are ordered based on their start position. Blue regions indicate gaps in the alignment which generally correspond to the position of introns.

### RNA splicing remains functional in the absence of SF3B3

Thus far, we had assessed the dynamic effects SF3B3 had on gene expression through our endogenous reporter, but we had not directly assessed Pol II activity by studying nascent transcription. To address this issue, we initially degraded SF3B3 and measured nascent transcription following A-485 treatment using transient transcriptome sequencing (TT-seq) to provide a direct quantification of newly produced transcripts^54^. The TT-seq data revealed clear separation by treatment condition and high replicate concordance (**Figure S3A**). These data also independently validated the transcriptional reactivation of *MYC* transcription (**Figure S3B**) confirming the utility of our approach to report on changes in endogenous gene expression. Notably, following stringent filtering, the TT-seq data revealed a set of 281 genes which like *MYC* were repressed following A-485 treatment but restored in the absence of SF3B3 (**Figure 2B & Figure S3C**). We found that these genes were highly expressed at steady state and are, on average, shorter than other expressed genes (**Figures S3D-E**). They possess similar numbers of exons but have an increased G/C content and reduced intron length (**Figures S3F-H)**. Gene set enrichment analyses were not discriminatory, instead showing that these genes encompass a broad range of cellular processes (**Figure S3I**).

Our TT-seq results along with the restored expression of the *MYC-mNG* reporter raised the intriguing prospect that splicing remains functional despite the loss of SF3B3. Consistent with this hypothesis, western blot analysis showed no difference in the size (kDa) of MYC-mNG protein produced in the presence or absence of SF3B3 (**Figure 2C**) suggesting it is likely derived from properly spliced transcripts despite SF3B3 degradation. Previous studies using Pladienolide B (PlaB), an inhibitor of SF3B1^55^, elicit widespread splicing defects and global transcriptional arrest^37^. In agreement with these findings, we found that cells treated with PlaB showed a decrease in MYC-mNG expression (**Figure 2D**) and were unable to restore MYC-mNG expression following SF3B3 degradation (**Figure S3J**). As PlaB may not completely inhibit SF3B1 or may induce off target effects, we endogenously tagged SF3B1 with dTAG in the MYC-mNG background (**Figure S3K**) to facilitate direct comparison of SF3B3 and SF3B1 degradation. The results were both striking and unexpected; whilst degradation of SF3B1 led to a rapid and complete loss in reporter gene expression, degradation of SF3B3 had only a minor effect (**Figures 2D-E**). These data, emphasise that although SF3B3 physically interacts with SF3B1 as part of the U2 snRNP spliceosome complex it plays a functionally distinct role in transcriptional control.

In further support of these findings, we observed marked intron inclusion following acute PlaB treatment or SF3B1 degradation which was not present following SF3B3 degradation (**Figure 2F**). To study the broader effects of SF3B3 loss at all genes we used 4-thiouridine (4sU) to label nascent transcripts in SF3B3-dTAG cells after 18 hours of dTAG^-V1^ or DMSO (vehicle) treatment. We then enriched full-length nascent transcripts and sequenced them using the Oxford Nanopore PromethION platform (**Figure 2G**). A global analysis of replicate experiments classifying all introns within major expressed gene isoforms as spliced (all overlapped introns spliced), partially spliced (some but not all possible introns spliced), or unspliced (no possible introns spliced), showed a minimal effect on splicing efficiency in SF3B3 degraded cells (**Figure 2H**). As an exemplar, comparable levels of splicing were noted in cells with or without SF3B3 at genes such as GAPDH which are highly transcribed and contain multiple introns of varying length (**Figure 2I**). Taken together, our data clearly separates the effects of SF3B1 and SF3B3 in splicing and transcription respectively. They demonstrate that SF3B3 loss is largely dispensable for ongoing pre-mRNA splicing but influences transcriptional activity to negate co-activator inhibition raising the intriguing prospect that a structural component of the U2 spliceosome complex may function primarily in transcriptional regulation.

### SF3B3 loss causes promoter proximal escape and reduced processivity of RNA Pol II

To study the differences in transcriptional regulation with greater granularity we measured nascent transcription using Precision run-on sequencing (PRO-seq)^56,57^ three hours after degradation of SF3B1 and SF3B3. By comparing the PRO-seq signal in SF3B1 or SF3B3 degraded cells to their respective controls we get a normalized appreciation of how each of these components of the U2 snRNP complex influence Pol II activity. These data revealed that, in contrast to SF3B3, the loss of SF3B1 leads to a profound increase in signal at the TSS with a marked depletion in the gene body consistent with a paused polymerase that is unable to productively elongate (**Figures 3A-C & Figure S3L**). To coordinate our global splicing and transcription analysis, we performed TT-Seq at the same time point as our nanopore analysis of splicing, which showed marked changes in gene expression (**Figure S3M**) characterized by increased transcription in the early gene body that gradually tapered and was reduced towards the 3’ end of the gene (**Figures 3D-E**). These findings raised the prospect that SF3B3 has a regulatory role in Pol II processivity.

**Figure 3.**
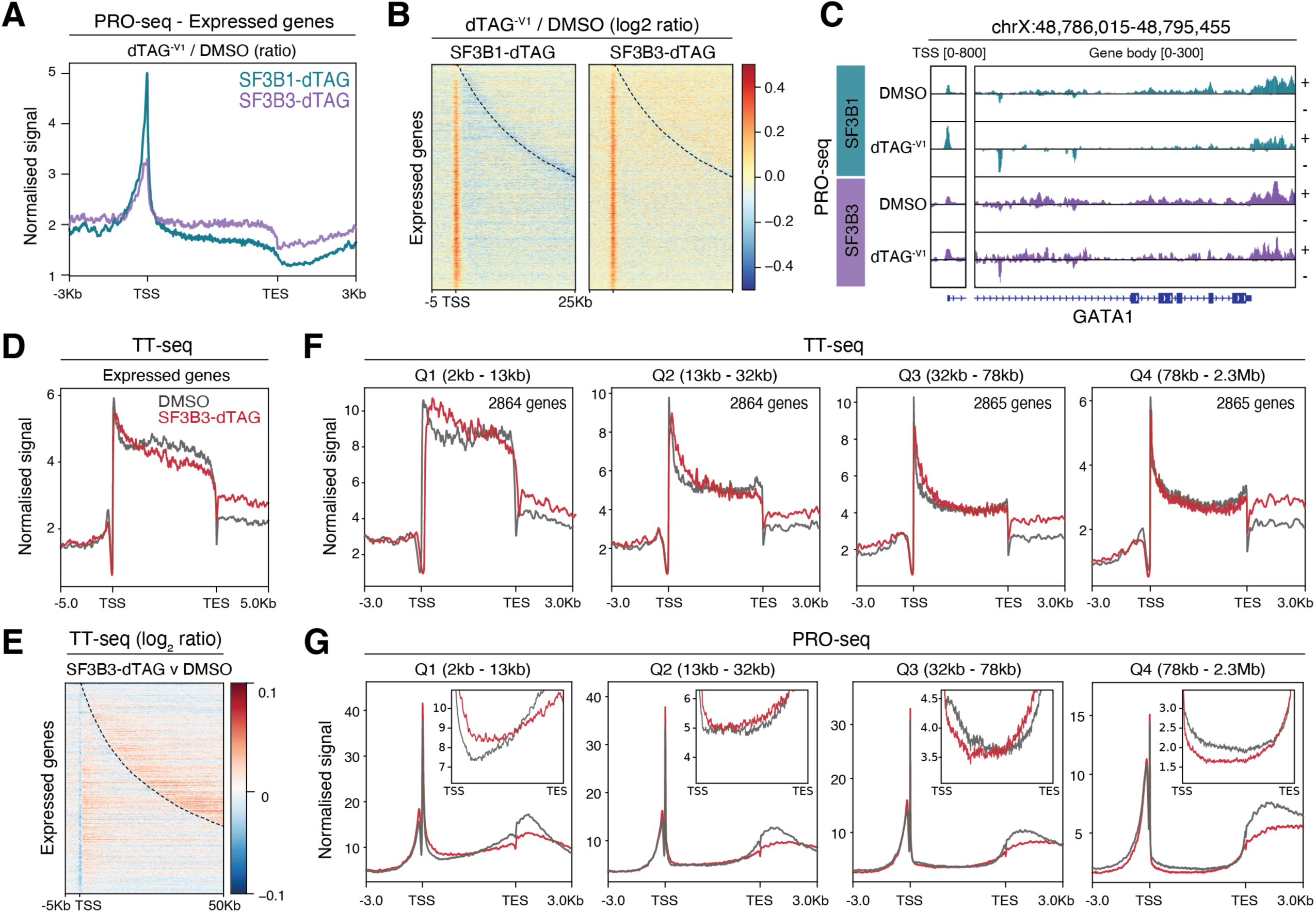
SF3B3 degradation causes a Pol II processivity defect. A) Metagene plot across all expressed genes (n = 12062) of PRO-seq experiments for SF3B1-dTAG and SF3B3-dTAG cells following 3h dTAG^-V1^ treatment. Profiles indicate ratio of dTAG^-V1^ / DMSO treatment conditions. Representative of n = 2 independent experiments. Scaled gene body +/- 5kb. TSS = transcription start site. TES = transcription end site. B) Heatmap of log_2_ ratio in PRO-seq signal for SF3B1-dTAG and SF3B3-dTAG vs. respective DMSO treatments. Heatmap ordered by ascending gene length. Dashed line indicates end of gene. TSS = Transcription start site. C) Genome browser visualisation of the *GATA1* locus for SF3B1-dTAG and SF3B3-dTAG PRO-seq data following 3h dTAG^-V1^ or DMSO treatment. Different scales used for TSS and gene body region. + /- indicates the strand orientation. Representative of n = 2 independent experiments. D) Metagene plot of TT-seq signal for all expressed genes in SF3B3-dTAG cells following 18h of DMSO or dTAG^-V1^ treatment (n = 12062). Scaled gene body +/- 5kb. Representative of n = 4 independent experiments. E) Heatmap of log_2_ fold change in TT-seq signal for SF3B3-dTAG vs. DMSO treatments at all expressed genes (−5kb to +50kb). Heatmap ordered by ascending gene length. Dashed line indicates end of gene. F) Metagene plots of TT-seq signal in SF3B3-dTAG cells following 18h of DMSO or dTAG^-V1^ treatment for expressed genes subsetted into Q1 (2-13kb), Q2 (13-32kb), Q3 (32-78kb) and Q4 (78kb-2.3Mb) gene length quartiles (n = 2864 genes / quartile). Scaled gene body +/- 5kb. G) Metagene plots of PRO-seq signal in SF3B3-dTAG cells following 18h of DMSO or dTAG^-V1^ treatment for expressed genes subsetted into Q1-Q4 subsets as in (f). Scaled gene body +/- 5kb. TSS = transcription start site. TES = transcription end site. Inset plots extend from TSS to TES with indicated y-axis limits.

To further investigate this, we apportioned expressed genes longer than 2kb into length-based quartiles (hereafter Q1-Q4 genes, each containing 2864 genes, **Table S5**), and) and analysed the transcriptional profile for each quartile. These data clearly showed that loss of SF3B3 leads to a marked increase in transcription immediately downstream of the TSS (**Figure 3F**). This finding, which was most pronounced at short genes, is consistent with impaired regulation of Pol II licensing and premature promoter proximal escape^58–61^. Importantly, these results also explain why the transcriptional restraint imposed by co-activator inhibition on short genes such as *MYC* is negated by the loss of SF3B3 (**Figure 2B, Figure S3E**). Notably, although the TT-seq data showed SF3B3 loss increased transcription early in gene bodies, particularly at Q1 and Q2 genes, they also reveal a marked tapering in transcription at the 3’end of the gene body (**Figure 3F**). To confirm these findings with a complimentary approach we generated further PRO-seq data which revealed a similar promoter proximal escape phenotype of Pol II, which was most pronounced at short genes along with a clear reduction in Pol II density at the 3’ end of the coding region, which is most marked at long genes (**Figure 3G & Figure S3N**). Together, these findings are most consistent with abnormal licensing of Pol II resulting in premature entry into elongation accompanied with a dramatic tapering in Pol II processivity.

### SF3B3 influences RNA Pol II pause release & elongation

To explain our nascent transcriptomics data, we surmised that SF3B3 degradation could impact Pol II activity in three major ways: 1) via differences in Pol II initiation; 2) via altered pause release efficiency; and/or 3) via changes in Pol II elongation kinetics following pause release. Focusing specifically on Q1 genes, which provided the greatest discriminatory opportunity, we examined total Pol II and Pol II Serine 5 CTD phosphorylation (pSer5) via ChIP-seq as a surrogate for Pol II initiation. We also assessed the chromatin occupancy of TATA binding protein (TBP) a key component of TFIID and the pre-initiation complex (PIC). These data show that SF3B3 degradation does not substantially alter the levels of either pSer5 or TBP at the TSS of these genes suggesting that SF3B3 does not significantly impact the assembly or initiation of the Pol II holoenzyme (**Figures 4A-C**). These findings were consistent when all expressed genes were examined suggesting this is a general feature of SF3B3 / Pol II dynamics rather than specific to the regulation of short genes (**Figures S4A-C**).

**Figure 4.**
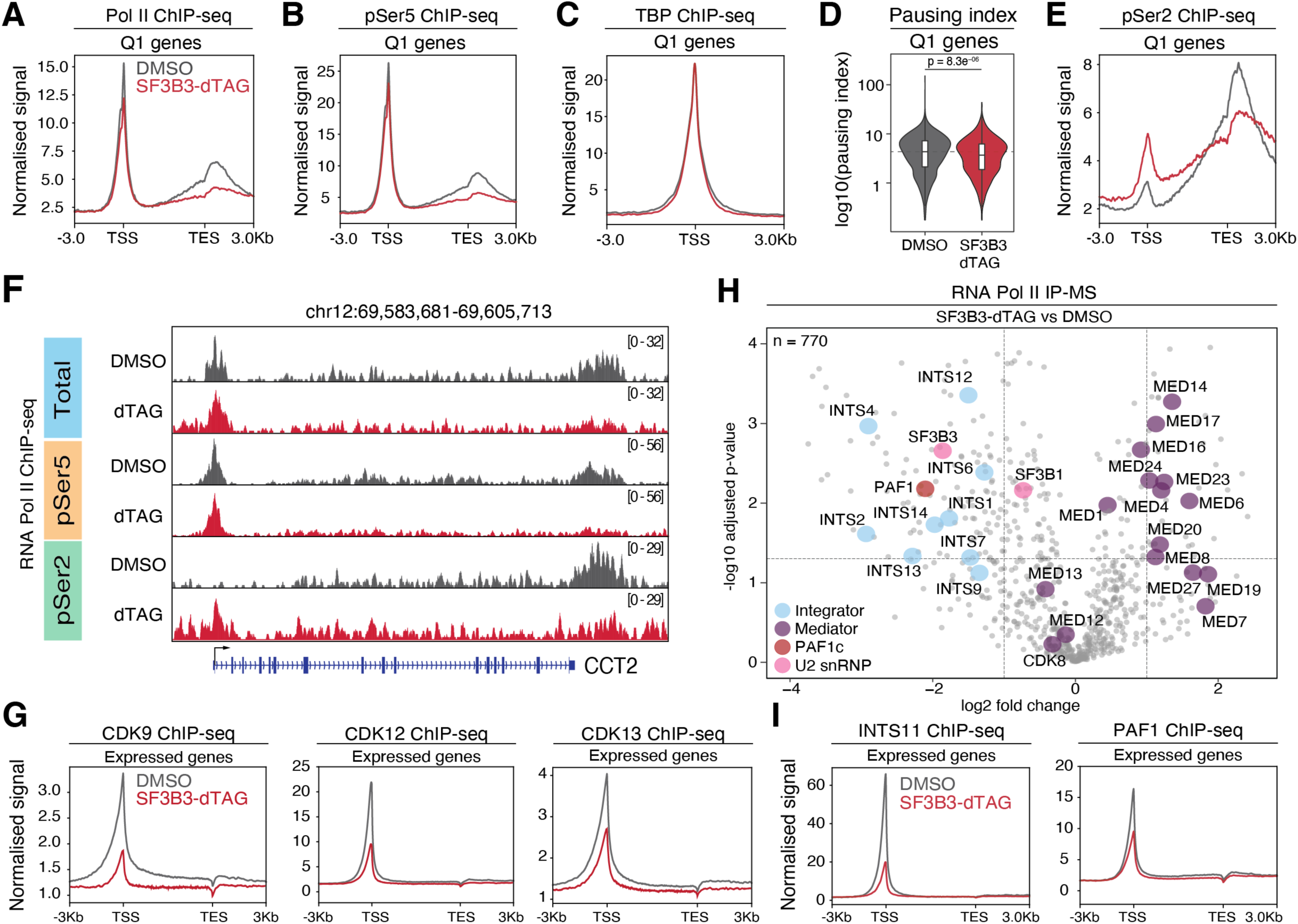
SF3B3 loss perturbs Pol II elongation via altered transcriptional cofactor and kinase recruitment. A) Metagene plot of total RNA Pol II ChIP-seq data at Q1 genes (n = 2864). Scaled gene body +/- 3kb. TSS = transcription start site. TES = transcription end site. Representative of n = 2 independent experiments. B) Metagene plot of Ser-5 phosphorylated RNA Pol II ChIP-seq data at Q1 genes (n = 2864). Scaled gene body +/- 3kb. Representative of n = 2 independent experiments. C) Profile plot of TATA binding protein (TBP) ChIP-seq data at Q1 genes (n = 2864). TSS +/- 3kb. D) Violin plot of total Pol II pausing index (promoter signal / gene body signal) at Q1 genes (n = 2864). log_10_ transformation. Two-sided Wilcoxon test vs. DMSO condition. Dashed line indicates the median of the DMSO condition. Boxplots span the upper quartile (upper limit), median (centre) and lower quartile (lower limit). Whiskers extend a maximum of 1.5x IQR. E) Metagene plot of Ser-2 phosphorylated RNA Pol II ChIP-seq data at Q1 genes (n = 2864). Scaled gene body +/- 3kb. Representative of n = 2 independent experiments. F) Genome browser visualisation of total Pol II, Ser-5 phosphorylated Pol II and Ser-2 phosphorylated Pol II ChIP-seq signal at the CCT2 locus in SF3B3-dTAG cells treated with DMSO or dTAG^-V1^. G) Metagene plots of CDK9, CDK12 and CDK13 ChIP-seq data at all expressed genes (n = 12062). Scaled gene body +/- 3kb. TSS = transcription start site. TES = transcription end site. Representative of n = 2 independent experiments. H) Volcano plot of RNA Pol II IP-MS data for dTAG^-V1^ (SF3B3-dTAG) vs DMSO treated SF3B3-dTAG cells. Dashed lines indicate an absolute log_2_ fold change threshold of 1 and an adjusted p-value threshold of 0.05 (-log_10_ transformation). Number of total proteins detected above background are shown (n = 770). Members of protein complexes of interest are indicated. I) Metagene plots of INTS11 and PAF1 ChIP-seq data at all expressed genes (n = 12062). Scaled gene body +/- 3kb. TSS = transcription start site. TES = transcription end site.

We next asked whether SF3B3 degradation alters Pol II pausing by calculating the Pol II pausing index, defined as the ratio of total Pol II ChIP-seq signal in the promoter proximal region (TSS −30bp to + 300bp) to the gene body (TSS + 300bp to Transcription End Site, TES)^62^. We found that degradation of SF3B3 significantly reduced the pausing index not only at Q1 genes but at all expressed genes (**Figure 4D & Figure S4D**). These findings suggest loss of SF3B3 leads to a reduced threshold to Pol II pause release, which is further supported by the increased early gene body transcription observed in our TT-seq and PRO-seq data. To explore this finding further we investigated the elongating form of Pol II via ChIP-seq for Pol II Serine 2 CTD phosphorylation (pSer2). Here we observed significantly increased pSer2 signal at the promoter proximal region of Q1 genes that dramatically tapered throughout the coding region and drops below control treatment levels towards the TES, mirroring our TT-seq and PRO-seq data (**Figure 4E & Figure S4E**). This trend was also observed at all expressed genes (**Figure S4F**). These findings illustrate the accumulation of elongating Pol II (**Figure 4F**) and are precisely the ChIP-seq pattern expected due to impaired Pol II processivity^63^. Together, our combined ChIP-seq, TT-seq and PRO-seq data shows that SF3B3 degradation has little impact on Pol II initiation but results in premature promoter escape and reduced processivity, collectively resulting in increased early gene body signal that rapidly diminishes towards the end of the gene body.

### SF3B3 coordinates the assembly of transcription elongation co-factors and cyclin-dependent kinases at gene promoters

Coordinated progression through the stages of transcription is dependent on the recruitment and catalytic activity of various cyclin dependent kinases (CDK). pSer2 is catalyzed by CDK9 as part of the active pTEFb complex and this is required for release of paused Pol II into productive elongation^64^. During elongation, pSer2 further accumulates on the Pol II CTD in a manner dependent on the transcriptional kinases CDK12 and CDK13, which facilitates the rate of Pol II elongation^27,65,66^. As our data showed marked abnormalities in the distribution of pSer2 at expressed genes in SF3B3 degraded cells, we sought to map the chromatin occupancy of CDK9, CDK12 and CDK13 using ChIP-seq. Our results showed that CDK9, CDK12 and CDK13 were primarily enriched at promoter proximal regions in vehicle treated cells which is consistent with their role in the modulation of Pol II activity (**Figure 4G**). However, following degradation of SF3B3 there was a marked reduction in the chromatin occupancy of all three CDKs that deposit pSer2 and facilitate Pol II elongation (**Figure 4G**).

Given the impact that SF3B3 loss has on genome-wide Pol II activity we also wanted to understand how SF3B3 influences the physical association of transcriptional complexes with Pol II. To address this question, we performed Pol II immunoprecipitation followed by label-free quantitative mass spectrometry (Pol II IP-MS), in the presence or absence of SF3B3. These data, generated from three independent biological replicates, were highly reproducible (**Figure S5A, Table S6**). In cells with intact SF3B3, relative to a bead only (mock) control, Pol II IP-MS enriched several protein complexes known to interact with Pol II including Mediator, SAGA, Integrator, PAF1 complex (PAF1c), and the U2 spliceosome with one of the top interacting proteins being SF3B3 (**Figure S5B**). When assessing the differential association with Pol II in cells with and without SF3B3, the Pol II interactome showed several notable changes, including an enhanced association with Mediator and a retained association with SF3B1. It is notable that SF3B1, the primary subunit, remains bound to Pol II even in the absence of SF3B3, explaining why splicing remains largely unaltered despite the loss of the third largest subunit of the U2 snRNP complex.

Conversely, following SF3B3 depletion we observed a significantly reduced interaction between Pol II and the Integrator and PAF1 complexes, suggesting that SF3B3 may coordinate the association of these complexes with the Pol II holoenzyme during transcription (**Figure 4H**). An impaired PAF1c-Pol II association is reported to result in premature promoter escape and reduced Pol II processivity during transcriptional elongation^58,67–69^, which is consistent with our TT-seq and PRO-seq findings after SF3B3 degradation (**Figures 3F-G**). Interestingly, many of the transcriptional consequences of SF3B3 loss are also phenocopied by impaired association of Integrator with Pol II. Disruption of Integrator association with Pol II, specifically the phosphatase module, would disrupt the dynamic equilibrium between elongation promoting kinases and the phosphatase PP2A which tempers this activity resulting in a reduced threshold for pause release^70–74^. Moreover, Integrator has also recently been shown to regulate the processivity of Pol II particularly in the region 12–35kb downstream of the TSS^61,75^. Therefore, whilst short genes are upregulated due to early Pol II release^73^, the impaired processivity in the absence of Integrator results in the downregulation of longer genes, which is similar to the phenotype we observe with SF3B3 loss. To validate our Pol II IP-MS findings at the chromatin template we performed ChIP-seq for INTS11, the catalytic subunit of the Integrator cleavage module^76^, and PAF1, a core subunit of PAF1c. These data confirmed the IP-MS findings, revealing that in the absence of SF3B3 there is a marked reduction in the chromatin occupancy of these key regulators that facilitate the elongation of Pol II (**Figure 4I**). Taken together, our findings illustrate that although SF3B3 is a structural component of the U2 spliceosome, it functions as a key factor that coordinates the recruitment and maintenance of Pol II transcriptional complexes, elongation factors and CDKs at chromatin. In its absence, an imbalance of key regulators of Pol II including CDK9, CDK12, CDK1313, Integrator and PAF1c arises, cumulatively resulting in promoter escape of Pol II and inefficient elongation (**Figure S5C**).

### The CTD of SF3B3 mediates the formation of the SF3B3/SF3B5 transcription module of the U2 snRNP complex

SF3B3, the third largest component of the SF3B complex^77^, consists of a three β-propeller (BP) WD40 domains (**Figure 5A**) that mediate interactions with splicing factors including the HEAT domain of SF3B1^78,79^. In addition to the BP domains, metazoan SF3B3 is predicted to contain a highly disordered C-terminal domain (CTD), whose molecular function is unclear (**Figure 5B**). The CTD includes a terminal stretch of 18 amino acids that is evolutionarily conserved only in metazoans (**Figure 5C**). To dissect the precise domain(s) of SF3B3 that contribute to the transcriptional phenotype, we ectopically expressed wild type SF3B3, or different truncation mutants of SF3B3 in the MYC-mNG reporter background (**Figure 5D**). We showed that each ectopic construct was expressed at comparable levels (**Figure S6A**). Next, we degraded endogenous SF3B3 using dTAG^-V1^ and examined the ability of each ectopic SF3B3 mutant to maintain transcriptional repression of the MYC-mNG endogenous reporter following coactivator inhibition.

**Figure 5.**
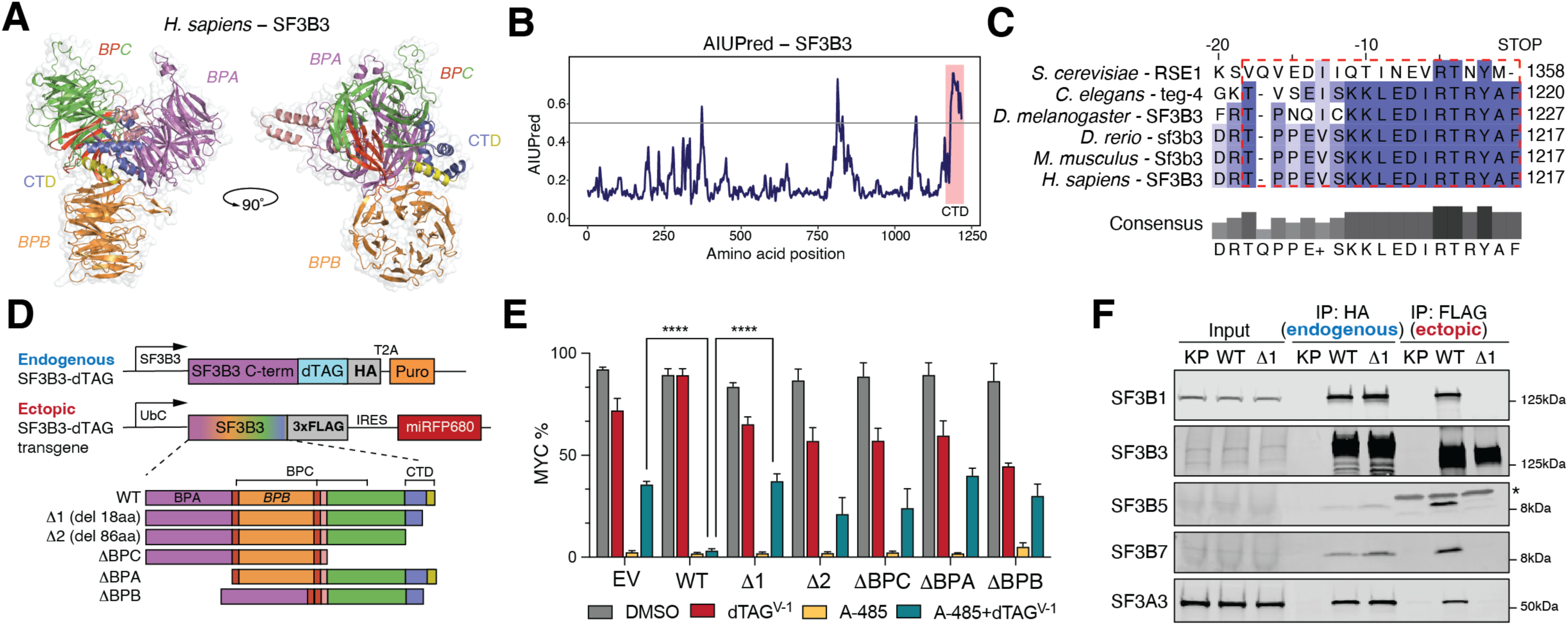
The terminal 18aa of SF3B3 C-terminal domain is metazoan specific and required for its effects on transcription. A) Structural prediction of *Homo sapiens* SF3B3 showing the locations of each b-propeller domain (BPA, BPB, BPC) and the C-terminal domain (CTD). B) AIUPred prediction of intrinsically disordered regions of *Homo sapiens* SF3B3. The default AIUPred score threshold of 0.5 was used. The SF3B3 CTD is highlighted. C) Protein sequence alignment of the final 20 amino acids of SF3B3 orthologs in unicellular and multicellular eukaryotes. Consensus score shown below alignment. Numbers indicate terminal amino acid position. CTD region deleted in the Δ1 mutant highlighted by the red box. D) Schematic of endogenous SF3B3-dTAG (HA-tagged) and the different ectopically introduced SF3B3 constructs (FLAG-tagged). E) Histogram of percentage MYC-mNG signal in MYC-mNG SF3B3-dTAG cells expressing the indicated SF3B3 overexpression construct pretreated with either 500nM dTAG^-V1^ or DMSO for 3 hours followed by treatment with either 2µM A-485 or 0.1% v/v DMSO for a further 20 hours. Percentage of MYC-mNG positive cells are indicated. Mean + standard deviation shown from n = 3 independent replicates. Two-sided Students T-test shown for indicated comparisons. F) Immunoblots following anti-HA (endogenous SF3B3) and anti-FLAG (ectopic SF3B3 construct) immunoprecipitation from cells expressing either wild type SF3B3 (WT) or the Δ1 SF3B3 mutant (Δ1). K562 parental cells used as negative control (KP). Representative of n = 3 independent experiments. * indicates a non-specific product in the SF3B5 immunoblot.

As expected, ectopic expression of wild type SF3B3 (WT) resulted in maintenance of transcriptional repression whist major structural alterations to SF3B3, such as deletion of each BP domain, phenocopied SF3B3 loss of function, resulting in transcriptional reactivation (**Figure 5E**). Surprisingly however, we found that truncation of just the CTD, particularly the terminal 18aa (Δ1), also phenocopied SF3B3 loss resulting in transcriptional reactivation following coactivator inhibition. To explore this result further we immunoprecipitated either WT SF3B3 or the Δ1 mutant and looked for association with other members of the U2 snRNP complex. These data revealed that the Δ1 mutant does not physically associate with several U2 snRNP complex members (**Figure 5F**) suggesting that the CTD enables the incorporation of SF3B3 into the U2 snRNP complex either directly, or via interaction with another protein.

To explore this possibility we next asked how the cellular proteome is impacted by acute SF3B3 perturbation by degrading SF3B3 and performing whole cell mass spectrometry four hours later. This revealed only one protein, SF3B5, which in the absence of SF3B3 was also significantly depleted (**Figure 6A**), suggesting that the interaction between these two proteins is required for their stability and incorporation into the U2 snRNP. Notably, SF3B5 was not one of the 1145 genes included in our chromatin focused sgRNA library that identified the transcriptional effects of SF3B3 (**Figures 1D-F**). Therefore, we generated an SF3B5-dTAG line to enable direct comparison with SF3B3 and SF3B1 (**Figure S6B**). These data revealed that loss of SF3B3 also results in the degradation of SF3B5 but not vice versa suggesting that SF3B5 needs to interact with SF3B3 for its stability. Interestingly, loss of SF3B1 does not destabilize the levels of either SF3B3 or SF3B5 suggesting that these two proteins have a linked physical and functional dependency that is independent of the primary subunit of the U2 snRNP complex. In fact, when we compared the effects of depleting SF3B1, SF3B3 or SF3B5 on the assembly and splicing function of the U2 snRNP we noted that degradation of SF3B1 led to the complete loss of association between the various constituents of the SF3B, SF3A and U2AF complexes that comprise the functional U2 snRNP complex (**Figure 6B**). However, loss of either SF3B3 or SF3B5 had minimal effect on the association of these U2 spliceosome complex members (**Figure 6B**). Consistent with this finding, intron retention at exemplar loci was markedly perturbed with SF3B1 degradation but similar to our findings with SF3B3, SF3B5 degradation had no dramatic effects on splicing (**Figure 6C**). Next, we tested if SF3B5 loss of function also alleviates the transcriptional repression induced by co-activator inhibition. Similar to our findings with SF3B3, we found that loss of SF3B5 also leads to reactivation of MYC-mNG (**Figure S6C**).

**Figure 6.**
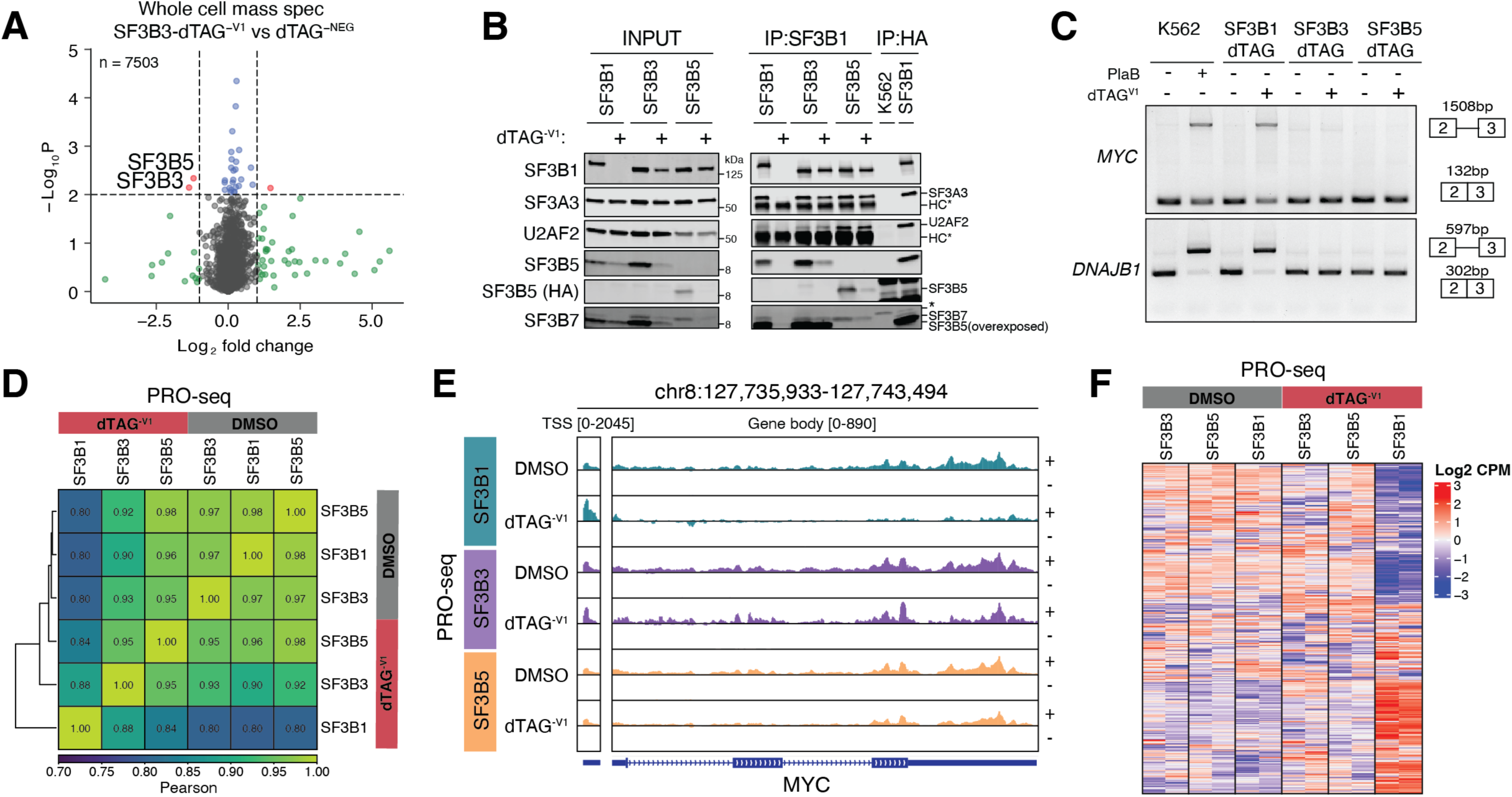
SF3B3 and SF3B5 form a metazoan specific U2 snRNP transcriptional submodule. A) Volcano plot of whole cell mass spectrometry data for SF3B3-dTAG cells treated with dTAG^-V1^ vs dTAG^-NEG^ for 3 hours. Dashed lines indicate an absolute log_2_ fold change threshold of 1 and an adjusted p-value threshold of 0.01 (-log_10_ transformation). Number of total proteins detected above background are shown (n = 7503). Proteins of interest are indicated. B) Immunoblots following anti-SF3B1 immunoprecipitation from SF3B1-dTAG, SF3B3-dTAG or SF3B5-dTAG cells treated with either dTAG^-V1^ or DMSO for 24 hours. Anti-HA (endogenous dTAG construct) immunoprecipitation from SF3B1-dTAG (positive control) or K562 parental cells (negative control, KP) used to control for non-specific and heavy chain products in the immunoprecipitated samples. Representative of n = 3 independent experiments. Non-specific and heavy chain products are indicated. C) Intron inclusion PCR assay across *c-MYC* exons 2-3 or *DNAJB1* exons 2-3 from MYC-mNG cells pretreated for 4 hours with 0.1% v/v DMSO or 100nM PlaB, or SF3B1-dTAG, SF3B3-dTAG or SF3B5-dTAG cells pretreated for 4 hours with 0.1% v/v DMSO or 500nM dTAG^-V1^. Schematic indicates expected size of spliced and unspliced amplicons. D) Pearson correlation of genome wide PRO-seq signal from SF3B1-dTAG, SF3B3-dTAG or SF3B5-dTAG cells pretreated for 3 hours with 0.1% v/v DMSO or 500nM dTAG^-V1^. Representative of n = 2 independent experiments. E) Genome browser visualisation of PRO-seq signal at the *MYC* locus in SF3B1-dTAG, SF3B3-dTAG and SF3B5-dTAG cells pretreated for 3 hours with 0.1% v/v DMSO or 500nM dTAG^-V1^. Representative of n = 2 independent experiments. F) Row-scaled heatmap of PRO-seq signal for SF3B1-dTAG, SF3B3-dTAG and SF3B5-dTAG cells pretreated for 3 hours with 0.1% v/v DMSO or 500nM dTAG^-V1^ at all expressed genes (n = 13587). Representative of n = 2 independent experiments.

To understand if the re-expression of MYC-mNG following SF3B5 depletion is also a consequence of premature promoter proximal escape of Pol II with impaired processivity we explored the transcriptional effects of acute SF3B5 degradation using PRO-seq. We found that the effects on Pol II activity following SF3B5 degradation were highly concordant with that of SF3B3 but distinct from the transcriptional effects seen with degradation of SF3B1 (**Figures 6D-F**). Interestingly, SF3B3 and SF3B5 have also been described to physically associate with metazoan SAGA (Spt-Ada-Gcn5 acetyltransferase), a highly conserved transcriptional co-activator complex primarily involved in transcriptional initiation and assembly of the pre-initiation complex^80–82^. Despite the physical association with SAGA in both plants and animals, the function of SF3B3 and SF3B5 within SAGA remains quite unclear^81^ with one study in *Drosophila* observing that loss of SF3B5 has both splicing abnormalities and transcriptional defects at some genes^83^. Although components of SAGA did not emerge in any of our functional screens, we wanted to understand if loss of some components of SAGA could phenocopy the transcriptional effects seen with loss of SF3B3 and SF3B5. In contrast to our findings with SF3B3/SF3B5 (**Figure S6C**), we found that knockout of several core and sub-module components of SAGA did not phenocopy the effects we observe with loss of SF3B3/SF3B5 (**Figure S6D**).

Taken together, our data show that the terminal 18aa of the unstructured SF3B3 CTD facilitates the interaction with SF3B5 which is needed for the stability of this submodule and proper incorporation into the U2 snRNP complex. This interaction is only present in metazoans, where remarkably this sub-module of the U2 snRNP complex does not dramatically impact splicing. Instead, it markedly influences the assembly and function of transcriptional kinases and co-activator complexes that enable optimal licensing and processivity of Pol II during transcriptional elongation.

## Discussion

Transcription of eukaryotic genes is a highly regulated multi-step process that is closely coordinated with co-transcriptional events including splicing. However, the reciprocal interplay between regulators of transcription and members of the spliceosome remains largely unclear. Studies investigating the role of splicing factors in modulating Pol II activity have largely relied on catalytic inhibition using Pladienolide B^37^ or snRNA anti-sense oligonucleotides^39^. In both cases, dramatic changes in splicing are observed, which concomitantly result in a striking disruption of transcription confirming that proficient splicing is necessary for efficient transcription. However, whether splicing factors can regulate transcription in a manner that is independent of their role in splicing has remained elusive. Here, we provide experimental evidence that SF3B3 and SF3B5, evolutionarily conserved members of the U2 spliceosome, play a major role in gene regulation by coordinating inputs from transcriptional co-activators and other co-factors to facilitate RNA Pol II pause release and elongation in a largely splicing-independent manner.

Through studying the endogenous regulation of *MYC*, we established that loss of SF3B3 results in promoter escape of Pol II, resulting in the enhanced expression of short, highly expressed genes, of which *MYC* is a prime example. Unfortunately, despite multiple approaches, we were unable to ChIP SF3B3 using several commercially available antibodies and/or epitope tags. Nevertheless, our IP-MS data confirmed a prominent association between Pol II and SF3B3^17,34^ and also established that the absence of SF3B3 results in a marked disruption to the equilibrium of Pol II elongation factors and transcriptional CDKs at chromatin. Consequently, the promoter escape of Pol II is accompanied by a dramatic decrease in Pol II processivity leading to the reduced expression of long genes (>30kb). SF3B3 is a large protein with three WD40 domains, a motif that are well established as a protein-protein interaction scaffold^84^. The association of SF3B3 with the Pol II CTD likely enables this scaffold function for the coordinated assembly of the transcriptional CDKs and elongation factors including PAF1c and Integrator. Notably, the WD40 domains are not the only important protein-protein interaction domains as the disordered CTD of SF3B3 is required for its association with SF3B5. The stability of this small 10 kDa protein is entirely contingent on the presence of SF3B3 and it plays a critical role in enabling the association of SF3B3 with the U2 snRNP. Although loss of SF3B1 results in the disassembly of the U2 snRNP complex, it does not alter the protein levels of SF3B3 or SF3B5 and conversely the loss of SF3B3 / SF3B5 does not affect the assembly or function of the U2 spliceosome nor does it perturb the association of SF3B1 with Pol II. These findings are consistent with the possibility that SF3B3/SF3B5 is a functionally distinct submodule, which although resident within the U2 snRNP complex, has co-evolved to coordinate transcription and splicing.

A broader question that warrants future consideration is why does a component of the spliceosome help coordinate Pol II promoter escape and processivity? Potentially, co-transcriptional splicing requires a precise rate of Pol II elongation for proper function and consequently we have evolved key components such as SF3B3 that exert a regulatory influence on Pol II to coordinate the efficiency of these processes. By convention, all structural components of a multi-subunit complex associated with a co-transcriptional process are ascribed a uniform function. For instance, all components of TREX are ascribed a role in nuclear export of mRNA and the components of the CPSF complex are linked by association with termination and polyadenylation. Our findings challenge this dogma and highlight the important point that structural components of a complex may have distinct regulatory functions. They raise the intriguing prospect that other multi-subunit complexes involved in other co-transcriptional processes may also contain structural components that function in a similar manner to SF3B3/SF3B5.

## Acknowledgements

We thank members of the Dawson laboratory for critical review of the manuscript and helpful advice. We also thank the Molecular Genomics Core, the Research Flow Core and the Victorian Centre for Functional Genomics (VCFG) at the Peter MacCallum Cancer Centre for helpful discussions and technical assistance, and Jafar Jabbari and Rory Bowden at the WEHI Advanced Genomics facility for assistance with Nanopore experiments.

## Author contributions

DV, JB & MAD conceptualised the research, designed and analysed experiments and wrote the manuscript with helpful contributions from all authors. DV & AG performed computational analyses. DV, JB, OB, WR, KP, OS and AD performed all experiments, analysed and interpreted data. CSA performed all mass spectrometry, generated and analysed data. DV and MAD jointly supervised the work.

## Data and materials availability

High throughput sequencing datasets have been deposited to the NCBI Gene Expression Omnibus under the GEO accession numbers GSE282773, GSE282779, GSE282782 and GSE301897. All code for analyses conducted in this study has been deposited at https://github.com/DaneVass/SF3B3_manuscript_2025.

## Supplemental information

Document S1. Figures S1-6

Table S1. Excel file containing sgRNA sequences for the bespoke human chromatin library

Table S2. Excel file containing oligos used in this study

Table S3. Excel file containing antibodies used in this study

Table S4. Excel file containing ChIP conditions used in this study

Table S5. Excel file containing expressed genes per length quartile group

Table S6. Excel file containing normalised label free quantification (LFQ) values from RNA Pol II IP-MS experiments

## Materials and Methods

### Compounds

I-BET151 was synthesised as previously described^47^. Nilotinib^48^ was obtained via materials transfer agreement with Novartis. Cycloheximide (cat #HY-12320), A-485^11^ (cat #HY-107455) and Pladienolide B^55,85^ (cat #HY-16399) were obtained from MedChemExpress. dTAG^-V1^ (cat #6914) and dTAG^-NEG^ ^52^ (cat #6915) were obtained from Tocris Bioscience.

### Cell lines and culture

*Homo sapiens* K562 erythro-leukaemia cells and *Drosophila melanogaster* S2 cells were obtained from the ATCC. *Homo sapiens* HEK-293ET cells were a gift from Dr. Felix Randow (MRC-LMB, Cambridge, UK). Human cell lines were authenticated by short tandem repeat profiling through the Australian Genome Research Facility. K562 cells were cultured at 37 °C and 5% CO_2_ in RPMI-1640 medium (Gibco #11875093) with 10% v/v heat inactivated fetal bovine serum (FBS), 100 IU/mL Penicillin, 100 g/mL Streptomycin and 2mM L-Glutamine (GlutaMAX, Gibco #35050061). HEK-293ET cells were cultured at 37 °C and 5% CO_2_ in DMEM (Gibco #10566016) with 10% v/v FBS, 100 IU/mL Penicillin, 100 g/mL Streptomycin., 2mM L-Glutamine (GlutaMAX). *Drosophila* S2 cells were cultured at 21 °C (room temperature) in Schneider’s *Drosophila* medium (Gibco #21720024) with 10% v/v FBS, 100 IU/mL Penicillin and 100 g/mL Streptomycin. Cell lines were routinely examined for Mycoplasma contamination by the Peter MacCallum Cancer Centre Genotyping core facility and were confirmed to be Mycoplasma negative throughout the duration of this study.

### Bespoke human chromatin CRISPR library generation

We designed a custom library of 7239 sgRNAs targeting 1146 genes encoding chromatin regulatory factors (6 sgRNAs per gene). The library included 186 non-targeting and 223 safe-harbor targeting control sgRNAs. The gene targets were identified based on protein domain content and subsequent manual curation (**Table S1**). sgRNA sequences were flanked by *BsmB*I cut sites and synthesized as an oligo pool by CustomArray (GenScript). Library sgRNAs were amplified by PCR and cloned into pKLV-U6gRNA(BbsI)-PGKpuro2ABFP (Addgene, #50946, a gift from Kosuke Yusa) which was modified to encode the human EF1a promoter in place of the human PGK promoter upstream of the eBFP-T2A-Puromycin cassette^43^. The ligated product was electroporated into 25µL Endura electrocompetent cells (Lucigen catalog #60242, 1.0 mm cuvette, 10 µF, 600 Ohms, 1800 Volts) using a GenePulser Xcell system (BioRad) and grown in Luria Broth overnight (∼16h) at 37 °C. The following day, the plasmid library was extracted via Maxiprep (Macherey-Nagel) and quantified via Qubit DNA broad range assay (Invitrogen). The composition and skew ratio (the ratio of the abundance of guide sequences in the top and bottom 10% of guide sequences^86^) of each library was determined by PCR amplification (see CRISPR/Cas9 screens section below) and high-throughput sequencing of the plasmid pool. Libraries with a skew ratio below 10 were considered of acceptable quality.

### Generation of K562 cMYC-mNeonGreen knock-in cell lines

To generate K562 cMYC-mNeonGreen (MYC-mNG) knock-in cell lines, a linear construct encoding the coding sequence of mNeonGreen^40^, T2A peptide, and a Blasticidin resistance cassette flanked by *c-MYC* sequences corresponding to the 500bp either side of the canonical stop codon was cloned into pUC19. Simultaneously, an sgRNA sequence targeting the 3’ terminus of the human *c-MYC* coding sequence (CDS) was cloned into pU6-(BbsI)_CBh-Cas9-T2A-mCherry (Addgene #64324, a gift from Ralf Kuehn) using *BsmB*I restriction digestion and ligation as described previously^87,88^. 300,000 parental K562 cells were suspended in Buffer R and electroporated with 500ng each of the sgRNA encoding vector and the MYC-mNeonGreen donor template encoding plasmid using the Neon Transfection System 10 µl kit (ThermoFisher, 1350V / 10ms / 4 pulses). After 48 hours of recovery post electroporation in culture medium without antibiotics, cells were transferred to antibiotic containing medium and selected with Blasticidin (20µg/mL) for 7 days. Individual blasticidin resistant, mNeonGreen positive cells were isolated using a FACS Aria II system into single wells of a 96 well plate and allowed to recover in medium containing 20µg/mL Blasticidin. Homozygous MYC-mNeonGreen positive clones were identified by PCR genotyping, flow cytometry and western blot for c-MYC. Sequences of sgRNAs, primers and donor templates are shown in **Table S2**.

### Lentivirus production and transduction

To prepare lentiviral vectors for small scale transduction experiments, 7×10^5^ HEK293ET cells in 2mL of media were plated per well in 6-well plates. The following day, 1000ng target vector, 350ng pMD2.G envelope vector (Addgene #12259, a gift from Didier Trono) and 700ng psPAX2 packaging vector (Addgene #12260, a gift from Didier Trono) were complexed with 6.15µg Polyethylenimine (PEI) in 100uL Opti-MEM (Gibco) and subsequently added to wells. Viral supernatant was collected at 48h and 72h following transfection, passed through a 0.45μm filter, and either used fresh or cryopreserved at −80°C. For routine transductions, 5×10^5^ K562 cells were incubated in 2mL of neat lentivirus overnight followed by media change. Unless stated otherwise, downstream assays commenced 48h-72h following transduction. For lentiviral preparations of bespoke CRISPR libraries, 17×10^6^ HEK293ET cells were plated per flask into at least 10 T175 flasks. Per flask, 17µg CRISPR library plasmid pool, 5.6µg pMD2.G envelope and 11.25µg psPAX2 packaging vectors were combined in a 3:2:1 weight ratio, complexed with 152µL (1mg/mL) Polyethylenimine (PEI) and added dropwise to the flask. Viral supernatant was collected 48h and 72h following transfection, combined, passed through a 0.45μm filter, aliquoted and cryopreserved at −80 °C.

### Generation of SF3B1, SF3B3 and SF3B5 dTAG knock-in cell lines

The SF3B1-dTAG, SF3B3-dTAG and SF3B5-dTAG knock-in cell lines were generated using the MMEJ/PITCh system as previously described^49^. Briefly, the hU6-PITCh-gRNA cassette from pX330S-2-PITCh (Addgene #63670) was subcloned into pX330A-1×2 (Addgene #58766), both gifts from Takashi Yamamoto, via *Bsa*I digestion to create a dual PITCh and target U6-sgRNA CRISPR–Cas9 vector labelled pX330-A+S. Forward and reverse DNA oligos encoding an sgRNA protospacer sequence at the N-terminus of SF3B1 or the C-terminus of SF3B3 or SF3B5 were annealed as described previously and ligated into pX330-A+S via *Bpi*I digestion and golden gate assembly^87^. Linear PITCh MMEJ dTAG-T2A-puromycin cassette (SF3B3-dTAG, SF3B5-dTAG) or PITCh MMEJ puromycin-T2A-dTAG cassette (SF3B1-dTAG) donor templates were generated by designing PCR oligos containing 20 bp of homologous sequence flanking the target locus. These were used in PCR reactions with pCRIS-PITChv2-dTAG-Puro (SF3B3-dTAG) or pCRIS-PITChv2-Puro-dTAG (Addgene #91796 and #91793; gifts from James Bradner & Behnam Nabet) as the template. Next, 750ng each of the matched sgRNA (pX330A+S) vector and donor template (pCRIS-PITChv2) were electroporated into 300,000 K562 MYC-mNG cells suspended in Buffer R using the Neon Transfection System 10 µl kit (ThermoFisher, 1350V / 10ms / 4 pulses). Cells were recovered for 48h in antibiotic free medium, followed by 5–7 days of puromycin selection (2µg/mL). Single-cell clones were isolated via FACS sorting into 96-well plates and allowed to expand in media containing puromycin. Following expansion, genomic DNA was isolated using DirectPCR Lysis Reagent (Viagen Biotech) according to the manufacturer’s instructions and used as input for genotyping PCRs. Clones demonstrating homozygous knock-in were further validated by Sanger sequencing of the homozygous knock-in gel extracted product and by immunoblot analysis using anti-SF3B1, anti-SF3B3, anti-SF3B5 and/or anti-HA antibodies. Oligonucleotide sequences used for dTAG cloning and PCR reactions are listed in **Table S2**.

### Timecourse proliferation and viability assays

50,000 cells were plated into 2mL total volume and treated as required. Daily volumetric counting was performed using a BD FACSverse (BD Biosciences). GFP signal (MYC-mNG expression) and cell viability (DAPI) was also recorded. Flow cytometry data were analysed with FlowJo v.10 (Tree Star).

### Antibodies

Antibodies used for immunoblot and/or ChIP assays are detailed in **Table S3**. The concentration used for each antibody is detailed in the relevant methods sections, and for ChIP assays in **Table S4**.

### Immunoblots

Cells were lysed in RIPA buffer (50 mM Tris (pH 7.4), 150 mM NaCl, 1 mM EDTA, 1% v/v Triton X-100, 0.5% v/v sodium deoxycholate, 0.5% v/v SDS supplemented with protease and phosphatase inhibitors (Roche). Lysate viscocity was reduced either by enzymatic (Benzonase; Merck) or mechanical (probe sonication) treatment. Normalized concentrations of protein were reduced and denatured for 5 min at 95 °C in Laemmli buffer containing either 10% v/v beta-mercaptoethanol or 100mM DTT and subsequently electrophoresed on 4–15% w/v precast polyacrylamide gels (Mini-PROTEAN TGX; Bio-Rad) under denaturing conditions. Proteins were wet-transferred onto methanol-activated PVDF membranes using the Mini Trans-Blot Electrophoretic Transfer Cell System (Bio-Rad) at 100 V (400 mA) for 60 min at 4 °C. Membranes were blocked for 1h at a room temperature of 22–25 °C in Intercept Blocking Buffer (LI-COR) and subsequently probed with primary antibody diluted at 1:1,000 (unless stated otherwise) in blocking buffer supplemented with 0.1% v/v Tween-20 overnight at 4 °C on a roller. Membranes were washed with TBS + 0.1% v/v Tween20 (TBST) and subsequently probed with the appropriate IRDye-conjugated secondary antibodies (LI-COR) diluted at 1:10,000 for 1h at room temperature. Membranes were scanned using an Odyssey Infrared Imaging System (LI-COR).

### Chromatin Immunoprecipitation (ChIP)

Approximately 25-30 ×10^6^ K562 cells were fixed with 1% v/v formaldehyde/PBS for 15 minutes (single crosslinking) or 2mM DSG/PBS for 30 minutes followed by the addition of 1% v/v formaldehyde for a further 15 minutes (dual crosslinking). Crosslinking was quenched with 250mM Tris pH 8.0. Crosslinked cells were lysed in shearing buffer (50mM Tris pH 8.0, 1% v/v SDS, 10mM EDTA supplemented with protease and phosphatase inhibitors). Chromatin was sonicated to approximately 250-1000bp fragments using the ME220 Focused-ultrasonicator (Covaris) and cellular debris was cleared by 20,000 rcf centrifugation for 15 min at 10 °C. Crosslinking and sonication conditions are further described in **Table S4**. Generally, 10-50µg of total sonicated chromatin was used for immunoprecipitation as quantified using the Qubit™ dsDNA quantification kit (ThermoFisher). Here, chromatin was diluted 10-fold with dilution buffer (55.55mM Tris pH 8.0, 166.65mM NaCl, 2.22mM EDTA, 1.11% v/v Triton X-100, supplemented with protease inhibitors) and subject to immunoprecipitation overnight at 4°C with rotation mixing. Antibodies used for ChIP are detailed in **Table S3**. Antibody bound complexes were immobilised with 50μL of pre-washed Protein A Dynabeads (Thermofisher) for 4 hours at 4°C with rotation mixing and collected using a magnetic rack. Beads were washed six times with wash buffer (50mM Tris pH 8.0, 150mM NaCl, 1% v/v Triton X-100, 0.1% v/v SDS, 2mM EDTA) followed by a final wash with TE buffer (10mM Tris pH 8.0, 1mM EDTA). Reverse crosslinking was achieved by incubating beads in elution buffer (20mM Tris pH 7.5, 200mM NaCl, 5mM EDTA, 1% v/v SDS, 200μg/mL Proteinase K (Roche) and 200µg/mL RNAse A (Qiagen)) at 65 °C from 4 hours to overnight with gentle agitation (1000 rpm). Immunoprecipitated DNA was purified using Qiagen MinElute columns (Qiagen). Sequencing libraries were prepared using the ThruPLEX® DNA-seq Kit (Takara Bio) as per manufacturer’s instructions and sequencing was subsequently performed on the Illumina NextSeq2000 with 100bp single-end chemistry targeting 30 million reads per sample.

### Intron inclusion PCR assays

Total RNA was column purified as per manufacturer’s instructions (RNeasy Kit; Qiagen). Isolated RNA was treated with DNase (Qiagen, cat #79254) to avoid genomic DNA contamination. cDNA was generated from 1µg of total RNA using random hexamer primers (Superscript VILO; ThermoFisher). PCR was performed on 1µL undiluted cDNA in a 50µL reaction using Q5® High-Fidelity DNA Polymerase (NEB) using the following thermocycling conditions: 120 seconds (s) at 98°C, 32 cycles of 20s at 98°C–10s at 62°C–15s at 72°C, and 5 minutes (min) at 72°C. A final concentration of 3% v/v DMSO was added to the PCR reaction for the *MYC* PCR. Amplified cDNA was resolved on a 2% w/v agarose gel. Primer pairs are listed in **Table S2**.

### Whole cell Proteomics

Cells were seeded at a density of approximately 3 × 10⁵ cells/mL and treated as specified in biological triplicates. After treatment, cells were washed twice with PBS and lysed in RIPA lysis buffer (as described above) supplemented with 250 U/mL Universal Nuclease (Pierce™) on ice for 30 minutes. Following clarification by centrifugation, protein concentration was determined using a BCA assay (Pierce™) and concentration normalised. SDS Buffer (9% SDS, 100 mM TEAB) was diluted 1.8-fold into the protein lysates to achieve a final SDS concentration of 5%, necessary for the S-Trap™ workflow. Protein digestion and purification were then performed using S-Trap™ Micro Columns (ProtiFi, LLC) according to the manufacturer’s instructions. Pooled eluates were freeze-dried overnight. Lyophilised peptides were resuspended in 2% (v/v) acetonitrile and 0.05% (v/v) Trifluoroacetic acid (TFA) and quantified using the micro BCA Protein Assay Kit (Pierce™). Each sample was measured in triplicate and normalised to a peptide concentration of 0.5 μg/mL prior to analysis by nanoESI-LC-MS/MS.

### Co-Immunoprecipitation of SF3B1, HA and FLAG

After acute 4-hour treatment with dTAG^-V1^, 1×10^7^ cells were subject to crude nuclear enrichment by resuspending cell pellets in ice-cold hypotonic lysis buffer (10mM Tris pH 8.0, 10mM NaCl, 5mM MgCl2, 0.2% v/v Tween 20 supplemented with fresh 0.02% w/v digitonin) for 10 minutes followed by centrifugation (800 rcf for 5 minutes). Following three washes with hypotonic buffer, nuclei were lysed in ice-cold lysis buffer (50mM Tris pH 8.0, 300mM NaCl, 1% Triton X-100, 1mM EDTA, 1mM EGTA) supplemented with protease inhibitors (Roche #11836170001) for 1 hour at 4°C with rotation-mixing. At the time of lysis, antibody-bead coupling was performed using an antibody to bead ratio of 5uL:30uL pre-washed protein A Dynabeads (Thermofisher) per sample. Antibody immobilisation was performed in TBST (TBS + 0.02% v/v Tween 20) at room-temperature with gentle agitation for 10 minutes and then stored on ice. Nuclear lysate was clarified by centrifugation (20,000 rcf for 15 minutes at 4°C) and subject to immunoprecipitation overnight at 4 degrees with mixing. The following day, co-immunoprecipitated material was washed 6 times with lysis buffer followed by 1 wash with TBS and proteins were eluted by boiling (95°C) for 5 minutes directly in 50uL of 1.2X Laemmli buffer. Note, to serve as negative controls for antibody heavy/light-chain contamination and to help further validate antibody specificity, parallel samples of untreated parental K562 and SF3B1-dTAG lines were processed identically, with the exception that magnetic anti-HA beads (Pierce™) were used for the immunoprecipitation.

### Co-Immunoprecipitation of RNA Pol II

RNA Pol II (POLR2A/RBP1) was immunoprecipitated as previously described with minor modifications^60^. Approximately 30 million cells were subject to crude nuclear enrichment by incubating cell pellets for 10 minutes in ice-cold hypotonic lysis buffer (10mM Tris pH 8.0, 10mM NaCl, 5mM MgCl_2_, 0.2% v/v Tween 20), supplemented with fresh 0.02% w/v digitonin, followed by centrifugation (800 rcf for 5 minutes at 4 °C). Following multiple washes with ice-cold hypotonic buffer, nuclei were lysed in 500µL of a chromatin digestion buffer (20 mM Tris-HCl pH 7.4, 150 mM NaCl, 1.5 mM MgCl_2_, 10% v/v Glycerol, 0.05% v/v IGEPAL, 0.5 mM DTT, protease inhibitors (Roche #11836170001), phosphatase inhibitors (Roche #4906845001) and ∼500 U/ml Benzonase (Sigma #E1014) for 1h at 4 °C with rotation mixing. Following centrifugation (20,000 rcf for 10 minutes), the supernatant was collected as fraction 1 and the residual chromatin pellet was subject to further extraction with 300µL of high-salt chromatin digestion buffer (20 mM Tris-HCl pH 7.4, 500 mM NaCl, 1.5 mM MgCl2, 10% v/v Glycerol, 0.05% v/v IGEPAL, 0.5 mM DTT, protease inhibitors, phosphatase inhibitors, 3 mM EDTA, 250U/mL Benzonase) with rotation mixing at 4 °C for a further hour. After centrifugation, the supernatant was collected, diluted with NaCl-free chromatin digestion buffer to normalise NaCl concentration to 150mM and finally pooled with fraction 1 to derive a total soluble nuclear extract. Pol II was immunoprecipitated from this extract using a monoclonal antibody against Pol II CTD (clone 4H8, Cell Signaling #2629S) crosslinked to Protein G Dynabeads (#10004D). Here, Dynabeads were washed and incubated with anti-Pol II antibody in PBST (PBS with 0.05% v/v Tween 20) (approximate ratio of 5µL Ab:40µL beads was used) for 4 hours at 4 °C with gentle rotation mixing. Uncoupled antibody was washed off and Dynabead-immobilised antibodies were crosslinked with 5mM BS3 (Thermo Scientific #A39266) in PBS for 30 minutes at room temperature with gentle agitation. Crosslinking reaction was immediately quenched by adding 1M Tris (pH 7.5) to a final concentration of 50mM and further incubated for 15 minutes. Crosslinked antibody-dynabead slurry was washed three times with chromatin digestion buffer and subsequently added to total nuclear extract for immunoprecipitation overnight at 4 °C with mixing. Co-immunoprecipitated Pol II complexes were washed 6 times in 1mL of wash buffer (20 mM Tris-HCl pH 7.4, 225 mM NaCl, 1.5 mM MgCl2, 10% v/v Glycerol, 1.5 mM EDTA, 0.05% v/v IGEPAL, 0.5 mM DTT) followed by a final wash in 100mM of TEAB (Thermo Scientific # 90114). Immunoprecipitated proteins were eluted in 25uL of 5% v/v SDS/8M Urea and subject to protein digestion and purification using S-Trap™ Micro Columns (Protifi) following manufacturer’s instructions. Pooled eluates were freeze-dried overnight. Lyophilised peptides were resuspended in 2% w/v Acetonitrile and 0.05% v/v Trifluoroacetic acid (TFA) and subject to nano-LC-MS/MS.

### LC-MS/MS and data analysis

Samples were analyzed by nanoESI-LC-MS/MS using an Orbitrap Exploris 480 mass spectrometer (Thermo Scientific) equipped with a nanoflow reversed-phase-HPLC (Ultimate 3000 RSLC, Dionex). The LC system was equipped with an Acclaim Pepmap nano-trap column (Dinoex-C18, 100 Å, 75 μm x 2 cm) and an Acclaim Pepmap RSLC analytical column (Dinoex-C18, 100 Å, 75 μm x 50 cm). The tryptic peptides were injected to the enrichment column at an isocratic flow of 5 μL/min of 2% v/v CH3CN containing 0.1% v/v formic acid for 5 min applied before the enrichment column was switched in-line with the analytical column. The eluents were 5% v/v DMSO in 0.1% v/v formic acid (solvent A) and 5% v/v DMSO in 100% v/v CH3CN and 0.1% v/v formic acid (solvent B). The flow gradient was (i) 0-6min at 3% B, (ii) 6-35 min, 3-23% B (iii) 35-45 min 23-40% B (iv) 45-50 min, 40-80% B (v) 50-55 min, maintain at 80% B (vi) 55-55.1, 80-3% B and equilibrated at 2% B for 10 minutes before the next sample injection. All spectra were acquired in positive ionization mode with full scan MS acquired from m/z 300-1600 in the FT mode at a mass resolving power of 120 000, after accumulating to an AGC target value of 3.0e6, with a maximum accumulation time of 25 ms. The RunStart EASY-IC lock internal lockmass was used. Data-dependent HCD MS/MS of charge states > 1 was performed using a 3 sec scan method, at a normalized AGC target of 100%, automatic injection, a normalized collision energy of 30%, and spectra acquired at a resolving power of 15000. Dynamic exclusion was used for 20 s.

Mass spectrometer raw files were processed in MaxQuant v2.6.2.0 with the Andromeda search engine and against the Uniprot Homo Sapiens FASTA sequences (downloaded June 2024). Default settings for protein and peptide identification were used. Cysteine carbamidomethyl was selected as fixed modification. Methionine oxidation and Protein (N-terminal acetylation) were set as variable modifications. The match between runs (MBR) and label free quantitation (LFQ) option was selected. Protein and peptides were filtered to a false discovery rate (FDR) of <0.01. MaxQuant output was imported into Perseus v2.0.11 for statistical analysis. Proteins that were labelled as “reverse”, and “contaminant” were removed from the dataset. The LFQ intensities were then log2-transformed and grouped according to the experimental groups. Proteins identified in all replicates for at least one experimental condition were retained for further analysis. Missing values were imputed (from a column normal distribution of 1.8 standard deviation down shift and 0.3 width) and normalised by median subtraction. To correct for multiple-hypothesis testing, significant hits are defined using Student’s t-test, truncated by permutation-based FDR threshold of 0.05 (250 randomization) and S0 factor of 0.1. Normalised LFQ data were exported and visualised using R v4.3.2. Normalised LFQ data is provided in **Table S6**.

### CRISPR/Cas9 gene disruption

Complimentary DNA oligonucleotides encoding an sgRNA targeting the gene of interest were synthesized by Integrated DNA Technologies. Oligos were annealed and cloned into the lentiviral sgRNA expression vector, pKLV-U6gRNA(BbsI)-PGKpuro2ABFP (Addgene, #50946, a gift from Kosuke Yusa), which was modified to encode the human EF1a promoter in place of the human PGK promoter^43^. Lentiviral vectors containing sgRNA encoding transgenes were produced as above. Cas9 expressing cells were transduced with an sgRNA of interest or an sgRNA targeting the human AAVS1 “safe-harbor” locus. Downstream assays were commenced 48h post sgRNA transduction to allow sufficient time for gene disruption to manifest. A list of sgRNA sequences is provided in **Table S2**.

### Generation of CRISPR/Cas9 expressing cell lines

For the temporal dependency and coactivator de-sensitisation screens, multiple clonal lines of MYC-mNG cells stably expressing a *S. pyogenes* Cas9-T2A-mCherry transgene were generated by transduction with lentivirus prepared from the FUCas9Cherry vector (Addgene #70182), a gift from Marco Herold, followed by single-cell flow sorting on mCherry expression (MYC-mNG Cas9-mCh). For reactivation dependency screens, multiple clonal lines of MYC-mNG SF3B3-dTAG cells stably expressing the *S. pyogenes* Cas9-T2A-mCherry transgene were generated using the same method. SpCas9 expressing clones were selected for further use based on robust and consistent mCherry expression by flow cytometry. All clonal populations of Cas9 expressing cells were subsequently validated for Cas9 expression and activity by flow cytometry and western blot prior to use in screens.

### CRISPR/Cas9 temporal dependency screens

The temporal dependency screens were conducted in biological quadruplicate over 6 days. At least 50×10^6^ MYC-mNG Cas9-mCh cells were transduced with human chromatin library virus at a multiplicity of infection of 0.2-0.3. Two days (48 hours) after transduction, viable mCherry (Cas9 reporter) and BFP (sgRNA reporter) double positive cells were sorted using a FACS Aria Fusion 3 or 5 system (BD Biosciences) and at least 8×10^6^ cells per replicate (equivalent to 1000x coverage of each library) were seeded into fresh media. On days 3,4,5 and 6 following transduction, at least 4×10^6^ cells were harvested from each replicate (library controls, minimum 500x coverage), resuspended in 200µL PBS and stored at −20 °C. Half of the remaining cells (at least 4×10^6^ cells) were reseeded in fresh media. mCherry (Cas9 reporter) positive, BFP (sgRNA reporter) positive and mNeonGreen negative (MYC reporter) cells were sorted from the remaining half (screen samples, at least 4×10^6^ cells) using a FACS Aria Fusion 3 or 5 system (BD Biosciences), resuspended in 200µL PBS and stored at −20 °C. On day 6 (the final day) no cells were reseeded and instead all remaining cells were sorted. To avoid cell confluency issues affecting MYC expression, media was refreshed on cells daily and cell cultures were maintained at 2.5-5×10^5^ cells/mL for the duration of the screen. Genomic DNA was isolated from library control and screen samples using the Qiagen DNeasy Blood and Tissue kit (Qiagen) with minor modifications. 5µL RNAse A (100mg/mL) was added to thawed cells in 200µL PBS and incubated at room temperature for 5 minutes before continuing with the rest of the protocol. gDNA was eluted in sterile water. Eluate was reapplied to the same column and incubated for a further 5 minutes before final isolation and quantification. Single guide RNA encoding sequences were amplified from all harvested gDNA using a one-step PCR approach with Q5 DNA polymerase (NEB) (see oligos in **Table S2**). We optimised the PCR reaction to amplify 1µg gDNA template over 28 cycles (98 °C – 3 minutes; 28 cycles: 98 °C – 30 seconds, 59 °C – 30 seconds, 72 °C – 30 seconds; 72 °C – 5 minutes). Approximately 10–20 PCR reactions were performed per screen sample dependent on the total amount of harvested gDNA. PCR reactions were pooled within each sample. 10µL was resolved on a 1.5% w/v agarose gel and the remainder was purified using the PCR and Gel cleanup kit (Machery Nagel) using manufacturer’s recommendations. Screen samples were sequenced on a NextSeq 500 using 75bp single end chemistry targeting 1×10^7^ reads for library controls and 5×10^6^ reads for screen samples.

### CRISPR/Cas9 coactivator de-sensitisation screens

The coactivator de-sensitisation screens were conducted in biological duplicate over 4 days. Nilotinib, A-485, I-BET151 and a DMSO control were screened in parallel. At least 50×10^6^ MYC-mNG Cas9-mCh cells were transduced and sorted after 48 hours as for the temporal dependency screens. At least 4×10^6^ cells per treatment and replicate (equivalent to 500x coverage of each library) were seeded into fresh media containing each drug (500nM Nilotinib, 2µM A-485, 2.5µM I-BET151). On day 4 following transduction (48h post drug addition), at least 4×10^6^ cells were harvested from each replicate (library controls, minimum 500x coverage), resuspended in 200µL PBS and stored at −20 °C. mCherry (Cas9 reporter) positive, BFP (sgRNA reporter) positive and mNeonGreen positive (MYC reporter) cells were sorted from the remaining cells (screen samples, at least 4×10^6^ cells), resuspended in 200µL PBS and stored at −20 °C. Genomic DNA extraction, sgRNA PCR and sequencing was conducted on library controls and screen samples as for temporal dependency screens.

### Flow cytometry

Fluorescence activated cell sorting was performed using a FACS Aria Fusion 3 or 5 flow sorter (BD Biosciences). Flow cytometry analysis was conducted on a Flow Symphony A3/A5, or Flow Fortessa (BD Biosciences). Flow cytometry data were analysed with FlowJo v.10 (Tree Star).

### Generation of SF3B3 overexpression constructs

The lentiviral vector expressing a C-terminal 3xFLAG-tagged SF3B3 was generated as follows: Originally an SF3B3-dTAG coding sequence was ordered as three separate gBlocks (IDT; **Table S2**) and stitched together alongside two separate PCR products encoding the miRFP680 cassette and IRES sequence using NEBuilder® HiFi DNA Assembly (NEB) as per manufacturer instructions. Fragments were assembled into a lentiviral backbone under the control of the human UbC promoter to minimise supraphysiological transgene expression. To generate the 3xFLAG version, the dTAG CDS was removed and replaced with a small gBlock encoding the C-terminal end of SF3B3 fused in-frame to a GSG-linker and the 3XFLAG peptide sequence using traditional digestion and ligation cloning. To generate the truncation mutants, back-to-back primers were designed to flank the region of interest for deletion. PCR amplicons were circularized using NEBuilder® HiFi DNA Assembly (NEB) as per manufacturer instructions. Mutagenesis primers are listed in **Table S2**. All vectors were verified by whole plasmid sequencing (Plasmidsaurus).

### High throughput 3’ RNA-seq (MAC-seq)

Multiplexed analysis of cells (MAC-seq) was performed as previously described^89^ with the following modifications. Total RNA was harvested from cells treated as required and 50ng was plated per well in 96 well plates. MAC-seq was performed on purified RNA omitting initial cell lysis steps. MAC-seq libraries were sequenced on a NextSeq 500 high output flow cell using 75bp cycle chemistry with the following cycle conditions: Read 1, 26 cycles; Index 1 read (i7), 6 cycles; Index 2 (i5), 0 cycles; Read 2, 60 cycles.

### Transient-transcriptome sequencing (TT-seq)

TT-seq was performed as described previously ^54^ with slight modifications. Briefly, Myc-mNG SF3B3-dTAG cells were pre-treated with either 500nM dTAG^-V1^ (Tocris Bioscience) or 500nM dTAG^NEG^ (Tocris Bioscience). After three hours, each group of pre-treated cells was exposed to either 2µM A-485 or 0.1% v/v DMSO (Vehicle control). 17.5 hours later, 4-thiouridine (4sU) was added to a final concentration of 100µM and cells were incubated for a further 30 minutes. Following this, 30 million cells from each condition were pelleted and resuspended in 5mL TRIzol reagent (Invitrogen). 2.4ng 4sU labelled Drosophila S2 total RNA was added to the TRIzol cell lysate per million cells prior to total RNA extraction, isopropanol precipitation and resuspension in sterile water. For RNA fragmentation, 300µg total RNA was diluted in 1mL cold 0.2M NaOH and incubated for 15 minutes on ice. The alkaline base hydrolysis reaction was quenched with 1mL 1M Tris-HCl pH 6.8. 300µg 4sU labelled total RNA was denatured at 65 °C for 10 minutes and placed on ice. Biotinylation was performed using EZ-link HPDP-Biotin (ThermoFisher Scientific) as previously described ^54^. 1µg of total 4sU labelled RNA was ethanol precipitated and kept aside for sequencing of total RNA abundance. Biotinylated nascent RNA was enriched from 300µg total 4sU labelled RNA using µMACS streptavidin beads (Miltenyi Biotec) as per the manufacturer’s recommendations. Isolated RNA was purified using the miRNeasy MinElute kit (Qiagen) with on-column DNaseI digestion performed as per the manufacturer’s recommendations. Sequencing libraries were generated from 50-200ng of total or nascent RNA fractions using the NEBNext® Ultra™ II Directional RNA Library Prep Kit for Illumina with ribosomal RNA depletion (New England Biolabs) and were sequenced on an Illumina NovaSeq S4 flow cell using 300 cycle chemistry (paired-end 150bp) targeting 30 million paired-end reads per sample.

### Precision Run-on sequencing (PRO-seq)

For PRO-seq assays 10 million MYC-mNG SF3B3-dTAG K562 cells were pre-treated with either 500nM dTAG^-V1^ or 500nM dTAG^NEG^. After three hours, each group of pre-treated cells were harvested and used as input for cell permeabilization. Alternatively, each group of pre-treated cells was exposed to either 2µM A-485 or 0.1% v/v DMSO (Vehicle control). 18 hours later, all cells from each condition were harvested and used as input for cell permeabilization. Aliquots containing one million permeabilized cells and 50,000 permeabilised *Drosophila* S2 cells were created per condition and used as input for the qPRO-seq assay^56,57^. Run on reactions with biotinylated dNTPs (PerkinElmer #NEL542001EA, #NEL543001EA, #NEL545001EA and #NEL544001EA) were performed for 5 minutes at 37 °C. Nascent RNA alkaline base hydrolysis was performed in ice cold 1M NaOH for 10 minutes. On bead reactions were performed as previously described^56^. Quantitative PCR reactions were performed to determine the number of cycles required for final library amplification. PRO-seq libraries were sequenced on an Illumina NextSeq2000 P2 flowcell using 100 cycle chemistry targeting 30 million reads per sample.

### Nascent long read RNA-sequencing using Oxford Nanopore

30 million MYC-mNG SF3B3-dTAG K562 cells were treated with either 500nM dTAG^-V1^ or 0.1% v/v DMSO. After 17.5 hours, 100µM 4sU was added and cells were incubated for a further 30 minutes. At the 18-hour mark cells were pelleted and resuspended in 5mL TRIzol reagent (Invitrogen) and total RNA was extracted, isopropanol precipitated and resuspended in sterile water. 300µg total RNA was biotinylated, enriched on beads, and purified as for TT-seq with DNaseI digestion performed on-column as per the manufacturer’s recommendations (Qiagen). 400 ng of total RNA was reverse transcribed using Maxima H Minus Reverse Transcriptase (ThermoFisher), following the protocol outlined in the PCR-cDNA Barcoding Kit (SQK-PCB111.24, Oxford Nanopore Technologies). Subsequently, 15 μL of cDNA was used for amplification with unique barcode primers in a 75 μL reaction. The amplification protocol included an initial denaturation at 95 °C for 30 seconds, followed by 12 cycles of 95 °C for 15 seconds, 62 °C for 15 seconds, and 65 °C for 8 minutes, with a final extension at 65 °C for 10 minutes. Post-amplification, Exonuclease I treatment was applied, and the resulting products were purified using 0.6x SPRIselect beads (Beckman Coulter) before being eluted in 20 μL of Nanopore Elution Buffer (EB). Equimolar amounts of the barcoded amplified cDNA were pooled to a final concentration of 25 fmol for the addition of Rapid Adapter T (RAP T). The final library was then loaded onto a PromethION FLO-PRO002 flow cell for sequencing.

### CRISPR knockout screen analysis

sgRNA counts were extracted from raw sequencing reads in fastq format, aligned to the reference sgRNA library and enumerated using BARtab v1.4 ^90^ with the following parameters: upconstant = CTTGTGGAAAGGACGAAACACCG, downconstant = “GTTTAAGAGCTATGCTGGAAAC”, constants = “both”, min_readlength = 16, alnmismatches = 1. Counts were imported into R v4.3.2 and differential sgRNA enrichment analysis was performed using edgeR v4 ^91^.

### ChIP-seq analysis

Quality of sequenced reads was assessed with FastQC v0.11.6 and adapters were trimmed with cutadapt v1.14^92^. Reads were then aligned to the Hg38 reference genome using Bowtie2 v2.3.4.1^93^ with the --no-unal flag in addition to default settings. Samples containing a Drosophila chromatin spike-in were also aligned to the BDGP6 reference genome in the same manner. Spike-in normalisation factors were derived using total *Drosophila* counts as previously described^94^. Duplicates were removed using the Picard toolkit v2.6.0. Sorting and indexing were performed with SAMtools v1.9^95^. Narrow and broad peaks were called with MACS2 v2.1.1^96^. Normalised bigwig files were generated using the bamCoverage function within Deeptools v3.5.5^97^. Normalisation was performed using 1x Reads per genomic content (1x RPGC) method or reference spike-in normalisation factor scaling where available. Heatmaps and metagene profiles were generated with Deeptools v3.5.5^97^. Read count plots per genomic regions were generated with BRGenomics v1.14.1 ^98^.

### 3’ RNA-seq analysis

Quality control and read trimming was performed as for ChIP-seq analysis. Reads were aligned to the hg38 genome and BDGP6 using STAR v2.7.5b^99^ with parameters --readFilesCommand zcat --outSAMtype BAM Unsorted --outFilterMultimapNmax 1 --outFilterMismatchNmax 3 in addition to default settings. Samtools v1.9^95^ was used for sorting and indexing of aligned BAM files. Picard v2.6.0 was used for removing duplicates from alignments. Read counts over genes were obtained with HTSeq v0.11.2^100^. Differential expression analysis was conducted using edgeR v4^91^. For 3’ MAC-seq data, STAR was run in STARsolo mode with the following parameters: --soloType CB_UMI_Simple --soloCBstart 1 --soloCBlen 10 --soloUMIstart 11 --soloUMIlen 10 --soloBarcodeReadLength 25 --outSAMattributes NH HI nM AS CR UR CB UB GX GN sS sQ sM --outSAMtype BAM SortedByCoordinate.

### PRO-seq analysis

Quality checking performed with FastQC v0.11.6. Adapter trimming was conducted using fastp v0.23.4^101^ using a custom adapter fasta file with additional parameters --umi --umi_loc=read1 --umi_len=6. Trimmed reads were aligned to Hg38 and BGDP6 reference genomes with Bowtie2 v2.3.4.1^93^ (with --local alignment setting in addition to default parameters). Compression to BAM format, quality filtering, sorting, and indexing was performed with Samtools v1.13^95^. Duplicates were removed with Picard v2.6.0. HTSeq v0.11.2^100^ was used for read quantification. Differential analysis was conducted using edgeR v4^91^. Genes were grouped into length-based quartiles using R v4.3.2.

### TT-seq analysis

Read QC and trimming was performed as for ChIPseq. Alignment and read counting performed as for RNAseq. Deduplication and BAM processing was conducted as for PROseq. Differential expression analysis was performed with edgeR v4^91^. Normalised bigwig files were generated using the bamCoverage function within Deeptools v3.5.5^97^. Normalisation was performed using 1x Reads per genomic content (1x RPGC) method or Drosophila reference spike-in normalisation factor scaling where available. Heatmaps and metagene profiles were generated with Deeptools v3.5.5. To define expressed genes, the TMM normalised^102^ total RNA-seq fraction from late timepoint TT-seq data were filtered to remove genes expressed below 0.5 TMM normalised counts in all samples. A BED file of expressed genes was generated by intersecting remaining Ensembl IDs post filtering with hg38 MANE gene annotations downloaded in GTF format from the UCSC table browser. To define the reactivated gene set the following steps were taken: (i) identify significantly downregulated genes in the A-485 condition vs DMSO (adjusted p < 0.05 & log_2_ fold change < 0); (ii) identify significantly upregulated genes in the SF3B3-dTAG condition vs DMSO (adjusted p < 0.05 & log_2_ fold change > 0) and filter these from the list of all expressed genes, leaving only genes that did not significantly increase in expression following SF3B3 degradation; (iii) identify genes common between lists (i) and (ii) that were significantly upregulated in the SF3B3-dTAG + A-485 condition vs A-485 (adjusted p < 0.05 & log_2_ fold change > 0). For length-based analyses, expressed genes longer than 2kb were grouped into length determined quartiles using R v4.3.2.

### Oxford Nanopore data analysis

Base calling was performed using MinKNOW software (version 23.4.6) with high-accuracy mode enabled. Data QC, alignment, and quantification was performed with nf-core/nanoseq pipeline v3.1.0 with Nextflow v23.04.1 and Singularity v3.7.3 with additional flags --skip_demultiplexing --skip_fusion_analysis --skip_modification_analysis --protocol cDNA --quantification_method stringtie2. Following this, secondary alignments were filtered with Samtools v1.17. Resulting BAM files were visualised in IGV v2.18. BAM files were read into R v4.3.2 using the GenomicAlignments package. Reads aligning to expressed genes were filtered and classified on their splicing status in R v4.3.2.

### SF3B3 disorder and evolutionary conservation analysis

Intrinsically disordered regions of *Homo sapiens* SF3B3 were predicted using AIUPred v0.1^103^. SF3B3 orthologous amino acid sequences from *Homo sapiens* (SF3B3), *Mus musculus* (Sf3b3), *Danio rerio* (sf3b3), *Drosophila melanogaster* (SF3B3), *Caenorhabditis elegans* (teg-4) and *Saccharomyces cerevisiae* (RSE1) were downloaded from OrthoDB v12.0 and aligned using CLUSTAL Omega v1.2.4. Alignments were visualised and annotated in Jalview v2.11.4.1.

---

**Figure S1.**
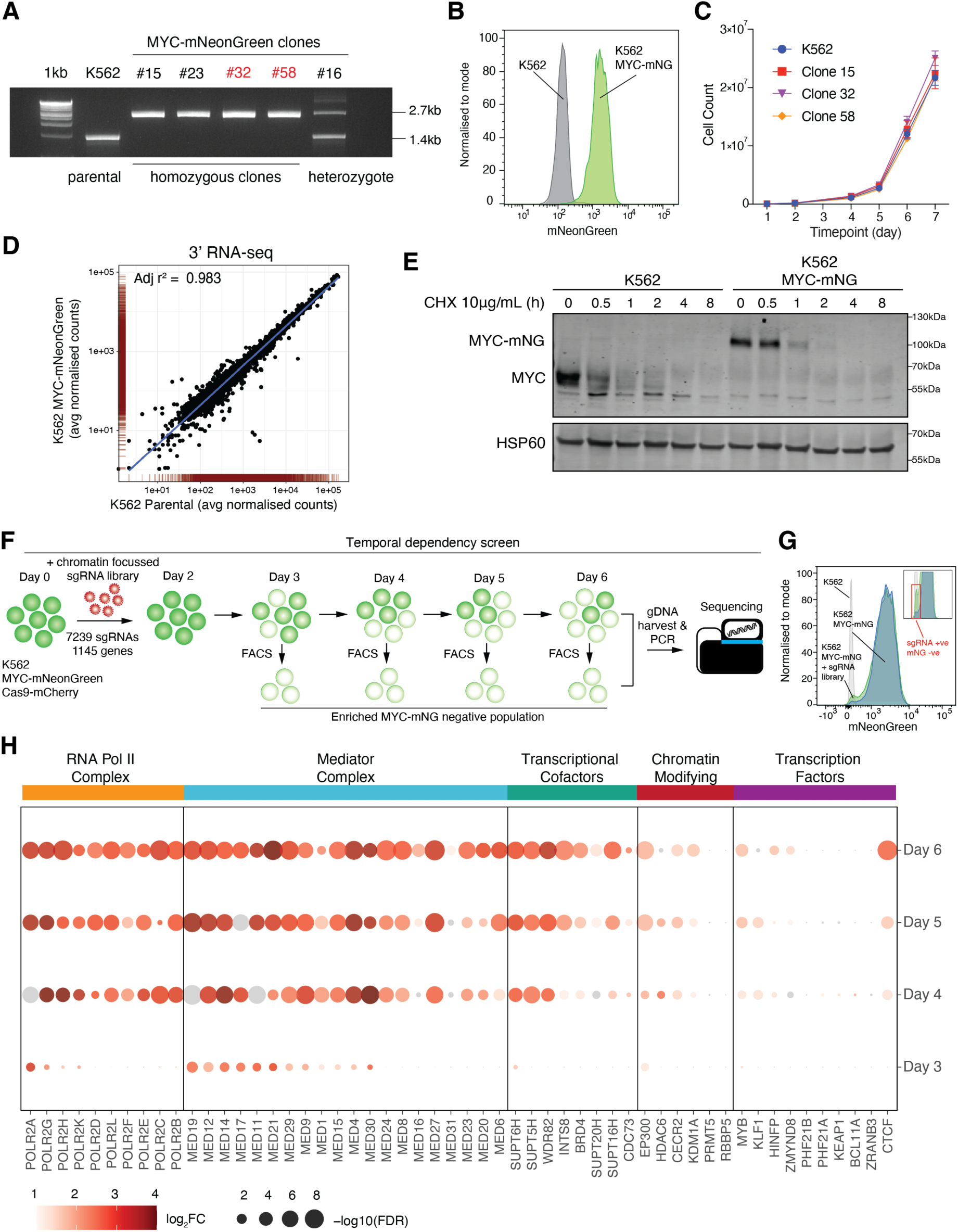
Temporal dependency screens using a real-time gene expression reporter system reveal varied contribution of transcriptional co-factors to gene expression. A) PCR validation of K562 MYC-mNG clonal populations. Homozygous clones #32 and #58 were used interchangeably throughout the study. B) Representative flow cytometry data (normalised to mode) for mNeonGreen in K562 parental cells (grey) and K562 MYC-mNG cells (green). C) Proliferation assay for K562 MYC-mNG clonal populations #15, #32, #58 and K562 parental cells. Mean + standard deviation shown from n = 3 independent replicates. D) Correlation of 3’ RNA-seq (MAC-seq) normalised counts for all expressed genes in K562 parental cells (average n = 4 replicates) vs. K562 MYC-mNG cells (average n = 2 replicates clone #32 and n = 2 replicates clone #58). Linear line of best fit (blue line) with adjusted r^2^ value as indicated. E) Western blot of c-MYC and HSP60 (loading control) following cycloheximide pulse chase assay in K562 parental and K562 MYC-mNG cells. Cells were dosed with 10µg/mL cycloheximide (CHX) for the indicated times (hours, h). Representative of n = 2 independent experiments. F) Experimental schematic for the CRISPR temporal dependency screen. G) Representative temporal dependency CRISPR screen FACS data indicating gating strategy to isolate reporter negative cells (inset). H) Dotplot of results for temporal dependency screens for the Day 3 to Day 6 timepoints (post sgRNA library transduction). Average of n = 4 independent experiments. Colour scale indicates log_2_ fold change (log_2_FC) enrichment in MYC-mNG negative population vs. control. Dot size indicates -log_10_ false discovery rate (FDR) corrected p-value. Grey points are significant based on -log_10_ adjusted p-value (FDR) < 0.05 but not log_2_FC > 1. Hits are categorised into RNA Pol II components (orange), Mediator components (blue), co-factors / co-activators (green), chromatin modifying enzymes (red) and transcription factors (purple).

**Figure S2.**
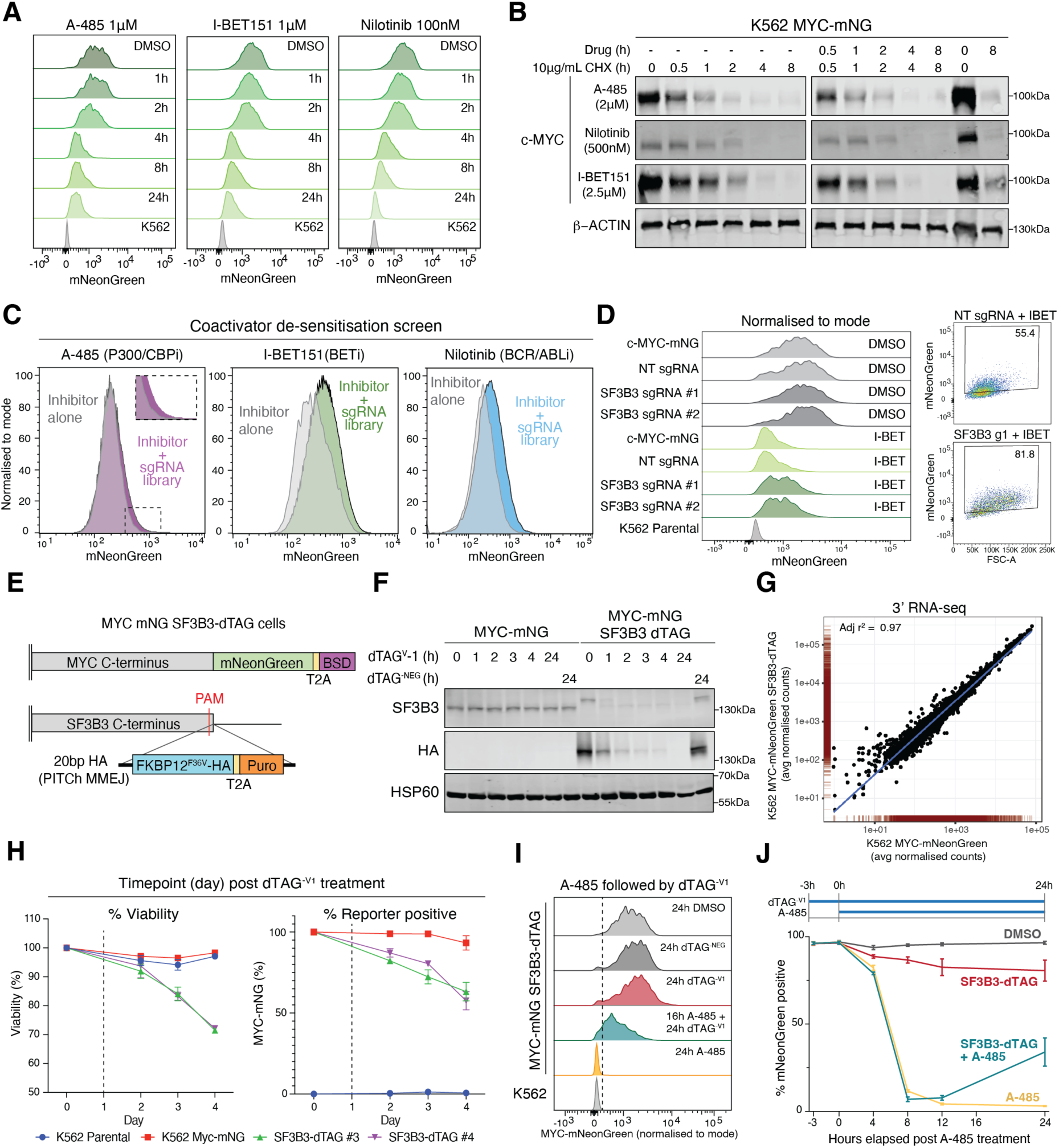
De-sensitisation screens reveal factors mediating sensitivity to transcriptional co-activator inhibition. A) mNeonGreen flow cytometry data (normalised to mode) in K562 MYC-mNG cells dosed for the indicated times (hours) with 1µM I-BET151, 100nM Nilotinib or 1µM A-485. DMSO (0.1% v/v) and K562 parental cells included as controls. Representative of n = 2 independent experiments. B) Western blot of c-MYC and β-ACTIN (loading control) following cycloheximide pulse chase assay in K562 MYC-mNG cells dosed with or without 2.5µM I-BET151, 500nM Nilotinib or 2µM A-485 for the indicated times (hours). Where indicated, cells were additionally dosed with 10µg/mL cycloheximide (CHX) for the times shown (hours). Representative of n = 2 independent experiments. C) Representative coactivator de-sensitisation CRISPR screen FACS data (normalised to mode) for Nilotinib (blue), A-485 (purple) or I-BET151 (green). Grey population indicates cells treated with each respective inhibitor only. D) Flow cytometry histograms (normalised to mode) and dotplots (right) following genetic knockout of SF3B3 in MYC-mNG cells with two independent sgRNAs and treatment with either DMSO (0.1% v/v) or 2.5µM I-BET151. Representative of n = 2 independent experiments. E) Schematic of dTAG (FKBP12F36V-HA-T2A-Puromycin) cassette knock-in into the endogenous SF3B3 locus at the C-terminus in K562 MYC-mNG cells. F) Western blot of SF3B3, HA (dTAG cassette) and HSP60 (loading control) following 500nM dTAG-V1 or 500nM dTAG-NEG treatment for the indicated times (hours, h) in MYC-mNG and MYC-mNG SF3B3-dTAG cells. Representative of n = 3 independent experiments. G) Correlation of 3’ RNA-seq (MAC-seq) normalised counts for all expressed genes in K562 MYC-mNG SF3B3-dTAG (average of n = 4 replicates) vs. K562 MYC-mNG cells (average of n = 4 replicates). Linear line of best fit (blue line) with adjusted r2 value as indicated. H) Percentage viability (DAPI negative) and percent MYC-mNG reporter positive for indicated cell lines across a four-day period post 500nM dTAG-V1 treatment. Mean + standard deviation shown from n = 3 independent replicates. Dashed lines indicate the latest timepoint (24 hours) at which degron-based assays in this study were conducted. I) Flow cytometry histograms (normalised to mode) of K562 MYC-mNG SF3B3-dTAG cells pretreated with DMSO 0.1% v/v or 2µM A-485 followed by 500nM dTAG-V1 or 500nM dTAG-NEG for the indicated times. Flow cytometry was performed at 24 hours post dTAG-V1/NEG treatment. Dashed line indicates the maximum mNeonGreen signal of the K562 parental population. J) Line plot of timecourse flow cytometry data showing percentage MYC-mNG reporter positive events per treatment condition. Cells were pretreated with 0.1% v/v DMSO or 500nM dTAG-V1 for 3 hours before further treatment with 2µM A-485 or 0.1% v/v DMSO. Mean + standard deviation shown from n = 3 independent replicates.

**Figure S3.**
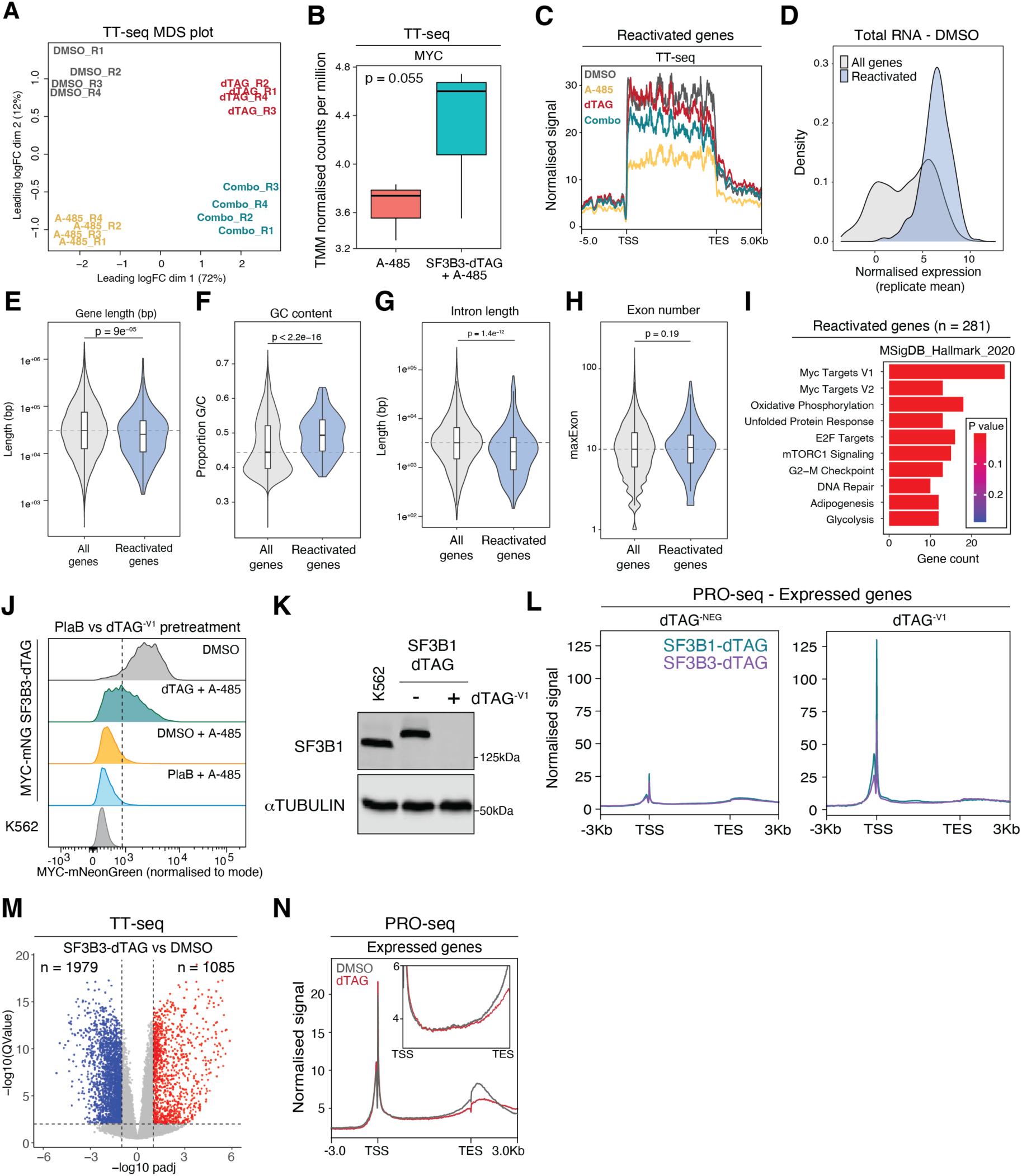
Reactivated genes are short and possess increased G/C content and reduced intron density. A) MDS plot of TT-seq (n = 4) data. B) Boxplots of TT-seq *MYC* expression data for A-485 and SF3B3-dTAG + A-485 treatments (N = 3 independent biological replicates). Two-sided students T-test. C) Metagene plot of TT-seq signal for reactivated genes (n = 281). Scaled gene body +/- 5kb. TSS = transcription start site. TES = transcription end site. Combo indicates SF3B3-dTAG + A-485 treatment. D) Density plot of normalised gene expression (averaged across n = 3 replicates) from total RNA-seq signal in the DMSO condition for all expressed genes (n = 12062) and reactivated genes (n = 281). E) Violin and boxplot of exon number for all expressed genes and reactivated genes. Two-sided Wilcoxon test. Dashed line indicates the median of all expressed genes. F) Violin and boxplot of proportion G/C content for all expressed genes and reactivated genes. Two-sided Wilcoxon test. Dashed line indicates the median of all expressed genes. G) Violin and boxplot of intron length (bp) for all expressed genes and reactivated genes. Two-sided Wilcoxon test. Dashed line indicates the median of all expressed genes. H) Violin and boxplot of exon number for all expressed genes and reactivated genes. Two-sided Wilcoxon test. Dashed line indicates the median of all expressed genes. I) MSigDB Hallmark 2020 gene set enrichment analysis for reactivated genes. J) Flow cytometry histograms of MYC-mNG expression (normalised to mode) in MYC-mNG SF3B3-dTAG cells pretreated with 500nM dTAG^-V1^ (dTAG), 100nM PlaB or 0.1% v/v DMSO for 3 hours followed by treatment with 2µM A-485 for a further 16 hours. K562 parental cells included as a negative control. MYC-mNG cells at steady state or after 16h 0.1% v/v DMSO treatment included as positive control. Dashed line indicates the maximum mNeonGreen signal of the K562 parental population. Representative of n = 3 independent experiments. K) Western blot of SF3B1 and alpha Tubulin (loading control) following 500nM dTAG^-V1^ or 0.1% v/v DMSO treatment for 24 hours in MYC-mNG SF3B1-dTAG cells. Parental K562 included as a control. Representative of n = 2 independent experiments. L) Metagene plot across all expressed genes (n = 12062) of PRO-seq experiments for SF3B1-dTAG and SF3B3-dTAG cells following 3h dTAG^-V1^ or 0.1% v/v DMSO treatment. Representative of n = 2 independent experiments. Scaled gene body +/- 5kb. TSS = transcription start site. TES = transcription end site. M) Volcano plot of TT-seq data for SF3B3-dTAG vs DMSO conditions. Dashed lines indicate absolute log_2_ fold change threshold of 1 and an adjusted p-value threshold of 0.01 (-log10 transformation). Number of significantly upregulated (red) and downregulated (blue) genes are shown. N) Metagene plot of PRO-seq signal for all expressed genes in SF3B3-dTAG cells treated for 18h with dTAG^-V1^ or 0.1% v/v DMSO. Scaled gene body +/- 5kb. TSS = transcription start site. TES = transcription end site. Inset plot extends from TSS to TES with y-axis limits 3-6. Boxplots span the upper quartile (upper limit), median (centre) and lower quartile (lower limit). Whiskers extend a maximum of 1.5x IQR.

**Figure S4.**
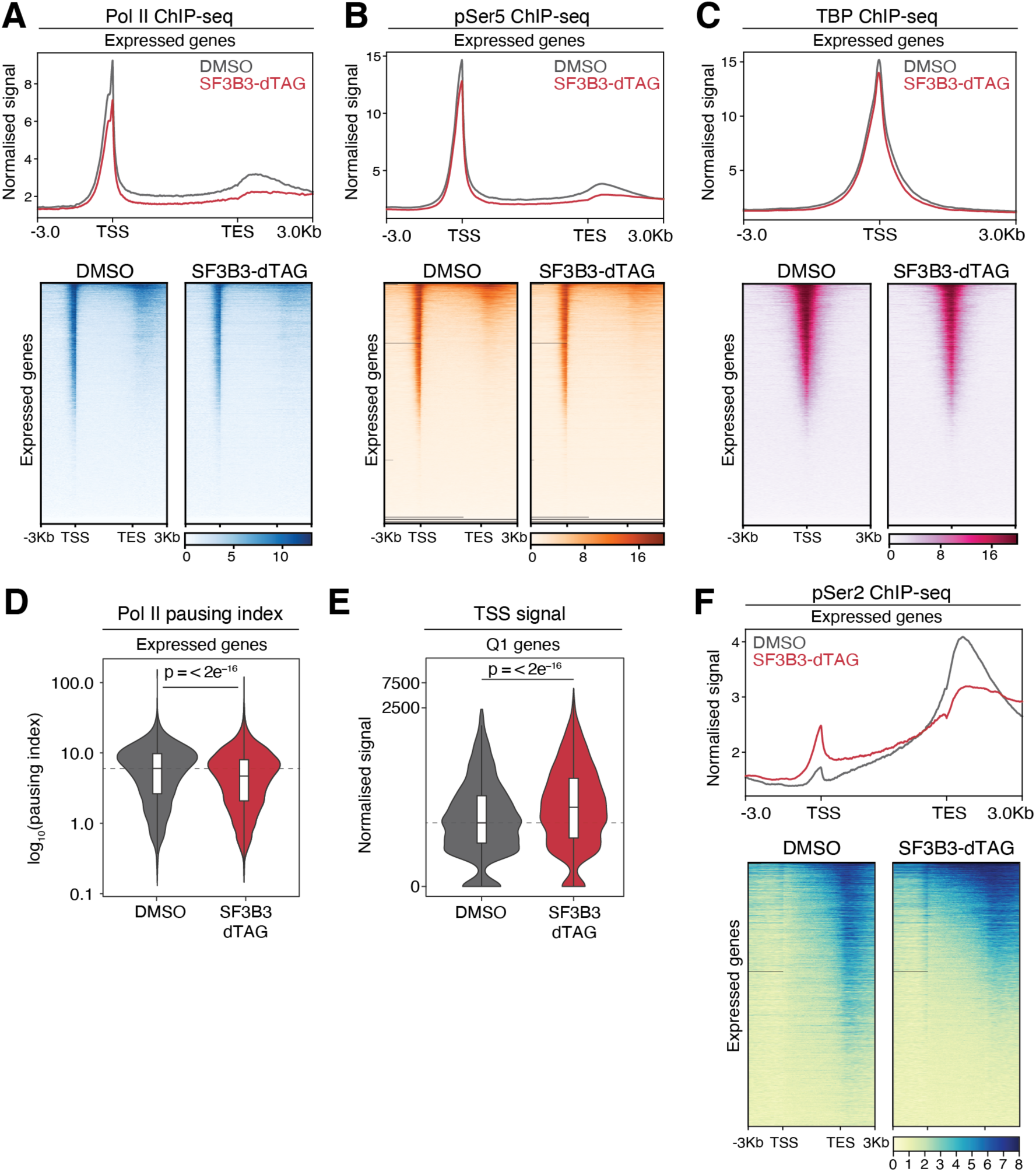
SF3B3 degradation perturbs RNA Pol II elongation. A) Metagene plot and heatmap of total RNA Pol II ChIP-seq data at all expressed genes (n = 12062) for SF3B3-dTAG cells treated with dTAG^-V1^ or DMSO. Scaled gene body +/- 3kb. TSS = transcription start site. TES = transcription end site. B) Metagene plot and heatmap of Ser5 phosphorylated RNA Pol II ChIP-seq data at all expressed genes (n = 12062) for SF3B3-dTAG cells treated with dTAG^-V1^ or DMSO. Scaled gene body +/- 3kb. TSS = transcription start site. TES = transcription end site. C) Profile plot and heatmap of TATA binding protein (TBP) ChIP-seq data at all expressed genes (n = 12062) for SF3B3-dTAG cells treated with dTAG^-V1^ or DMSO. TSS +/- 3kb. D) Violin and boxplots of total Pol II pausing index (promoter signal / gene body signal) all expressed genes (n = 12062) for SF3B3-dTAG cells treated with dTAG^-V1^ or DMSO. log_10_ transformation. Two-sided Wilcoxon test versus DMSO condition. Dashed line indicates the median of the DMSO condition. E) Violin plot of Ser-2 phosphorylated RNA Pol II ChIP-seq signal at the transcription start site (TSS, +/- 200bp) of Q1 genes (n = 2864) for SF3B3-dTAG cells treated with dTAG^-V1^ or DMSO. log1p transformation. Two-sided Wilcoxon test versus DMSO condition. Dashed line indicates the median of the DMSO condition. F) Metagene plot and heatmap of Ser2 phosphorylated RNA Pol II ChIP-seq data at all expressed genes (n = 12062) for SF3B3-dTAG cells treated with dTAG^-V1^ or DMSO. Scaled gene body +/- 3kb. TSS = transcription start site. TES = transcription end site. Boxplots span the upper quartile (upper limit), median (centre) and lower quartile (lower limit). Whiskers extend a maximum of 1.5x IQR.

**Figure S5.**
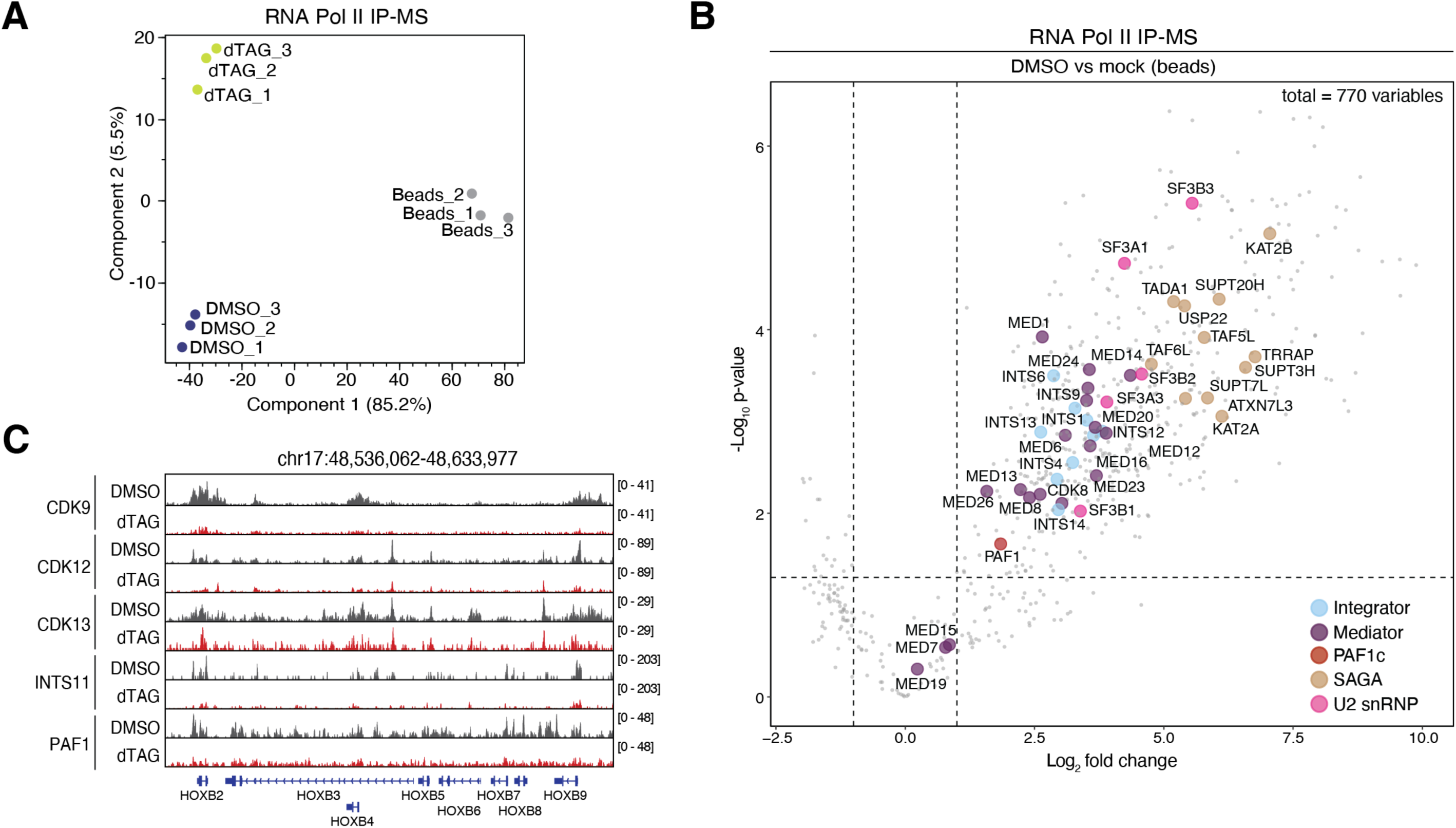
SF3B3 coordinates the assembly of transcriptional complexes with the RNA Pol II holoenzyme. A) Principal components analysis of RNA Pol II co-immunoprecipitation mass-spectrometry (IP-MS) data for SF3B3-dTAG cells treated for 24h with 0.1% v/v DMSO, 500nM dTAG^-V1^ or a mock (beads only) control. N = 3 independent experiments. B) Volcano plot of protein enrichment from RNA Pol II co-immunoprecipitation mass-spectrometry (IP-MS) data for SF3B3-dTAG cells treated for 24h with 0.1% v/v DMSO versus a mock (beads only) control. Dashed lines indicate an absolute log_2_ fold change threshold of 1 and an adjusted p-value threshold of 0.05 (-log_10_ transformation). Number of total proteins detected above background are shown (n = 770). Members of protein complexes of interest are indicated. C) Genome browser visualisation of CDK9, CDK12, CDK13, INTS11 and PAF1 ChIP-seq data at the *HOXB locus* for SF3B3-dTAG cells treated with dTAG^-V1^ or DMSO.

**Figure S6.**
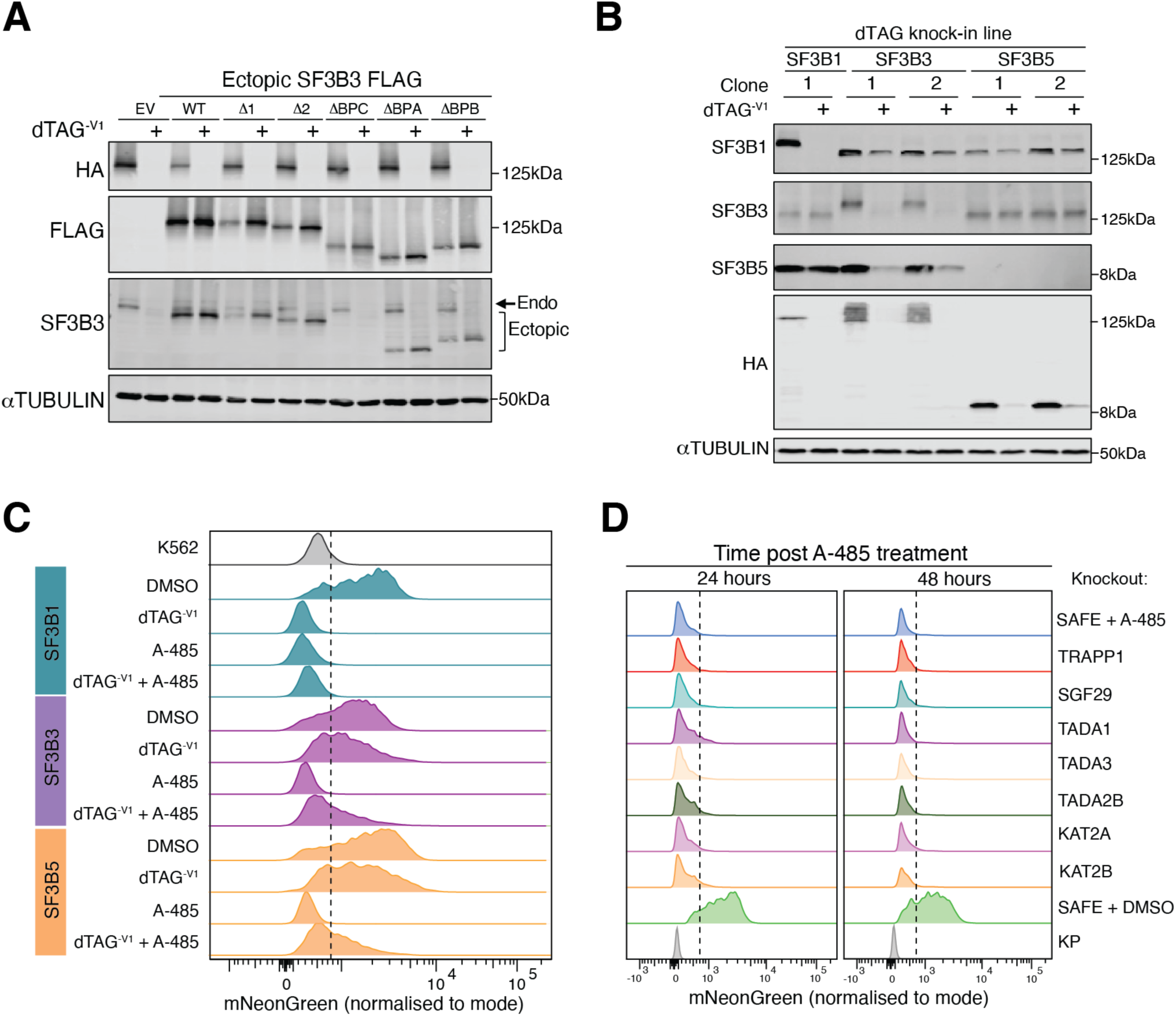
The C-terminus of SF3B3 modulates its association with SF3B5. A) Immunoblot showing expression levels of ectopic SF3B3 overexpression constructs in the SF3B3-dTAG background. Cells were treated with 500nM dTAG^-V1^ or 0.1% v/v DMSO for 24h. Endogenous SF3B3-dTAG is tagged with HA. Ectopic SF3B3 constructs are FLAG tagged. For the SF3B3 immunoblot, endogenous and ectopic isoforms are indicated. Representative of n = 2 independent experiments. B) Immunoblot of SF3B1 (n = 1 clones), SF3B3 (n = 2 clones), SF3B5 (n = 2 clones), HA (dTAG alleles) and alpha-Tubulin (loading control) for K562 parental cells (negative control) and the indicated endogenously tagged dTAG lines. Cells were treated with 500nM dTAG^-V1^ or 0.1% v/v DMSO for 24h. dTAG allele inhibits SF3B5 signal in the SF3B5-dTAG background. In these cells SF3B5-dTAG expression is shown by HA blot. Representative of n = 3 independent experiments. C) Flow cytometry histograms of MYC-mNG expression (mNeonGreen, normalised to mode) in SF3B1-dTAG (green), SF3B3-dTAG (purple) or SF3B5-dTAG cells (orange) treated with 0.1% v/v DMSO or 500nM dTAG^-V1^ for 3h followed by either 0.1% v/v DMSO or 2μM A-485 for 24h. Representative of n = 2 independent experiments. Dashed line indicates the upper signal limit of cells treated with A-485 alone. D) Flow cytometry histograms of MYC-mNG expression (mNeonGreen, normalised to mode) in MYC-mNG Cas9-mCH cells transduced with different constructs encoding sgRNAs against SAGA complex family members. 48 hours after sgRNA transduction, cells were treated with 2μM A-485 and mNeonGreen signal was assessed after 24 and 48h. SAFE indicates a control sgRNA targeting the *AAVS1* safe-harbor locus. SAFE + DMSO indicates cells transduced with a control sgRNA and treated with 0.1% v/v DMSO (negative control). Representative of n = 2 independent experiments.

